# Eco-evolutionary interaction in competing phytoplankton: genotype sorting likely explains dominance shift and species responses to CO_2_

**DOI:** 10.1101/2020.03.31.017954

**Authors:** Luisa Listmann, Giannina S. I. Hattich, Birte Matthiessen, Thorsten B.H. Reusch

## Abstract

How ecological and evolutionary processes interact and together determine species and community responses to climate change is poorly understood. We studied long-term dynamics (over approximately 200 asexual generations) in two phytoplankton species, a coccolithophore (*Emiliania huxleyi*) and a diatom (*Chaetoceros affinis*), to increased CO_2_ growing alone or competing with one another in co-occurrence. To allow for rapid evolutionary responses, the experiment started with a standing genetic variation of nine genotypes in each of the species. Under co-occurrence of both species, we observed a dominance shift from *C. affinis* to *E. huxleyi* after about 120 generations in both CO_2_ treatments, but more pronounced under high CO_2_. Associated with this shift, we only found weak adaptation to high CO_2_ in the diatom and none in the coccolithophore in terms of species’ growth rates. In addition, no adaptation to interspecific competition could be observed by comparing the single to the two-species treatments in reciprocal assays, regardless of the CO_2_ treatment. Nevertheless, highly reproducible genotype sorting left only one genotype remaining for each of the species among all treatments. This strong evolutionary selection coincided with the dominance shift from *C. affinis* to *E. huxleyi*. Since all other conditions were kept constant over time, the most parsimonious explanation for the dominance shift is that the strong evolutionary selection potentially altered competitive ability of the two species. Thus, here observed changes in the simplest possible two-species phytoplankton “community” demonstrated that eco-evolutionary interactions can be critical for predicting community responses to climate change in rapidly dividing organisms such as phytoplankton.

## 1 Introduction

Recent studies have repeatedly shown that ecological and evolutionary processes happen on similar time scales (Carroll et al., 2007; Reznick, 2013). Understanding how both processes interact has sparked the emergence of the new field of eco-evolutionary dynamics (Fussmann et al., 2007) aiming to understand how rapid evolutionary adaptation influences ecological processes, and vice versa (Hendry, 2016). Strong eco-evolutionary coupling is particularly expected in phytoplankton species, photoautotrophic aquatic microbes that can form large populations. With one cell division per day, many species lend themselves for experimental studies allowing for appropriate replication and hundreds of generations of evolutionary change (Reusch and Boyd, 2013). Moreover, their ecological importance cannot be overemphasized, as marine phytoplankton species are responsible for half of all global photosynthesis (Falkowski et al., 2008).

Here we studied eco-evolutionary dynamics in response to increased seawater CO_2_ concentration, one prominent aspect of ocean global change (Doney et al., 2009). As primary producers, all phytoplankton species depend on the availability of inorganic carbon. On time scales too short for evolutionary adaptation, non-calcifying species mostly respond by enhanced growth (Li et al., 2017; Schaum et al., 2012) leading to higher abundances (Sommer et al., 2015) (“ecological winners”, e.g. diatoms), while most calcifying species react negatively (“ecological losers”, e.g. coccolithophores) to CO_2_ enrichment (Doney et al., 2009; Riebesell, 2004). Nested within such broad functional categories of sensitivity is pronounced intraspecific variation (Des Roches et al., 2018; Hattich et al., 2017; Schaum et al., 2012), on which selection can operate (Lohbeck et al., 2012).

Within phytoplankton, competition for abiotic resources is ubiquitous (Tilman, 1977). In recent theoretical studies (de Mazancourt et al., 2008; Lancaster et al., 2017) and a long term bacterial community experiment (Lawrence et al., 2012), competition was shown to constitute one major biotic driver of adaptation. At the same time experimental evolution studies using phytoplankton and addressing the presence and absence of competition as a selection factor are largely absent (Scheinin et al., 2015).

Here, we set out to study how the evolutionary response to increased CO_2_ of two bloom forming and geographically co-occurring phytoplankton species, *Emiliania huxleyi* (a coccolithophore) and *Chaetoceros affinis* (a diatom), was altered by competition and assessed putative effects on ecological dynamics. Studying such eco-evolutionary processes requires that at least two species can be kept over long-term (i.e. more than 10s of generations such that evolutionary change can happen) in co-occurring experimental settings, delaying or preventing Gause’s principle of competitive exclusion (Hardin Garret, 1960; Tilman, 1977). Hence, we established long-term coexistence in semi-continuous batch cycles by taking advantage of the species’ different nutrient uptake related strategies (Doney et al., 2009; Riebesell, 2004; Sommer, 1984). Diatoms have high nutrient uptake rates and are consequently characterized by high maximum growth rates (Litchman et al., 2007). At the same time, diatoms have high half saturation constants and low affinity for nutrient uptake (Litchman et al., 2007). As such they represent the ‘velocity-adapted’ species (Sommer, 1984) and thrive in nutrient replete and fluctuating conditions, being able to rapidly monopolize nutrients. Coccolithophores in contrast, have lower maximum nutrient uptake rates. Hence, after a nutrient pulse they initially lose out against diatoms. Their lower half-saturation constants and higher affinity, however, make them the better competitors under nutrient poor conditions thriving at later successional stages (Sommer, 1984). In this system, we ran the experiment for ca. 200 asexual generations under fully factorial selection conditions of adaptation to ambient and increased CO_2_, in combination with and without competition, i.e. the co-occurrence of the respective other species (Fig. 1 a). To allow for rapid evolutionary adaptation via genotype sorting (Lohbeck et al., 2012), populations of both species were assembled using nine genotypes in equal frequencies (Fig. 1 a) that could be traced via microsatellite genotyping. Reciprocal adaptation assays were conducted to test for adaptation of both species to enhanced CO_2_ and the presence of the respective other species in the first and second half of the experiment (after ca. 50 and 200 generations) (Fig. 1 b).

**Figure 1.**
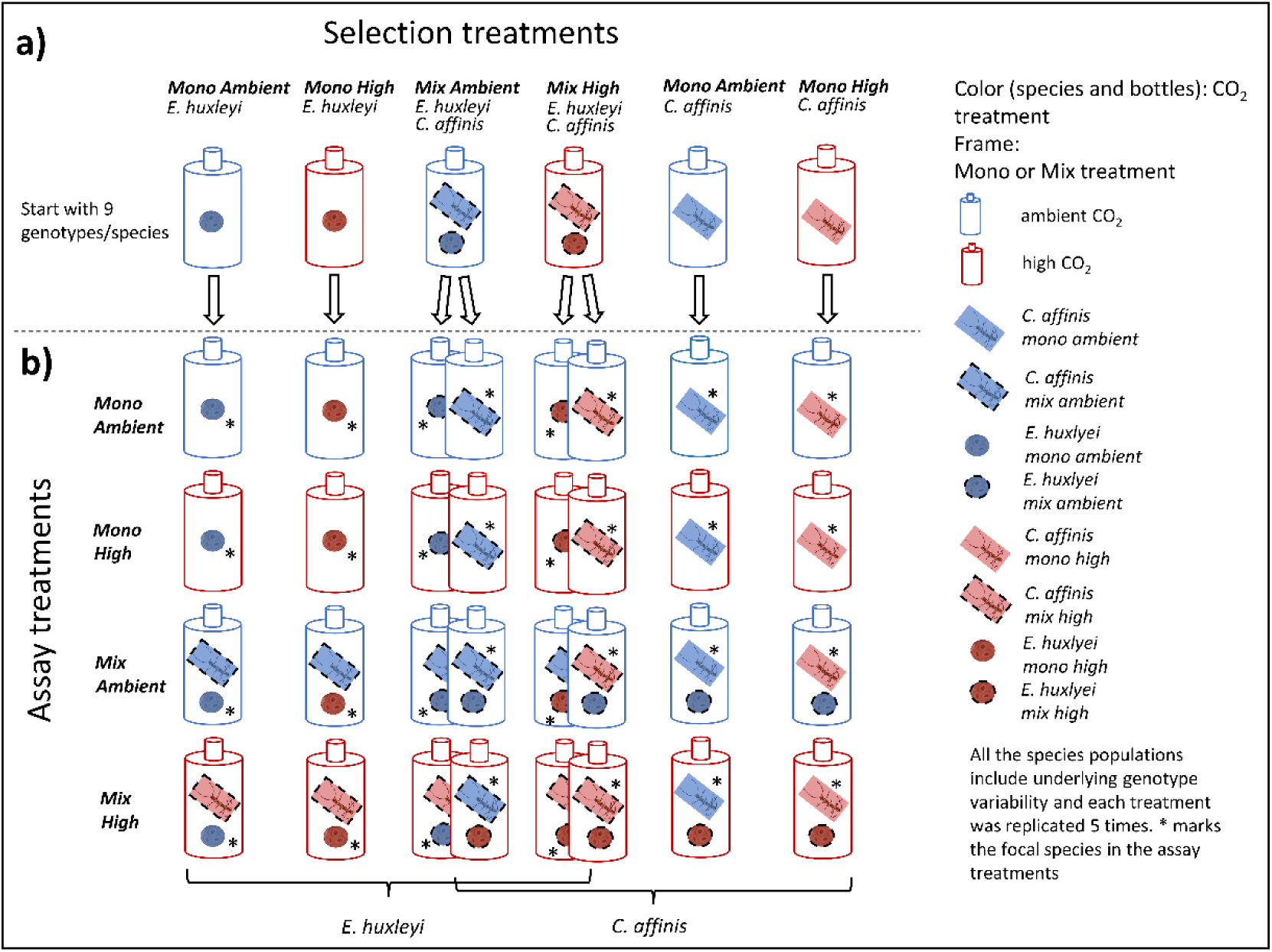
Set up of Selection and Assay experiments: Here the schematic setup of the selection phase of the experiment is depicted in the top row (a). The colors show the CO_2_ treatments, whereas the frames of the species show mono or mix treatment. All species populations were started with 9 genotypes each. Each treatment consisted of 5 replicates. In the reciprocal assays, all cultures of both species were exposed to all possible treatments (four rows below) thus resulting in a fully crossed assay set-up (b). The 16 resulting assay treatments are shown for each species. The star shows which of the species was the focal species in the mix assay treatments.

We hypothesize that on the short term, species contribution to the “community” would be diverging between CO_2_ treatments due to varying CO_2_ responses of the target species. Specifically, the relative contribution of *E. huxleyi* should be lower in the high CO_2_ compared to the ambient treatment due to potential negative or neutral effects of CO_2_. In contrast, the relative contribution of *C. affinis* should be higher in high CO_2_ treatment due to a potential fertilizing effect of increased CO_2_ in combination with the effect of CO_2_ on *E. huxleyi*. In the long-term, we hypothesize that evolutionary dynamics (i.e. genotype sorting and eventually adaptation) diverge between both target species. Specifically, the potentially negatively affected coccolithophore should be selected towards more stress tolerance, while the possible fertilizing effect on the diatom could lead to selection for the diatom’s potential to efficiently utilize increased levels of CO_2_ and in turn achieve higher growth rates. Moreover, any evolutionary change can feed back onto ecological processes by altering the abundance and thus contribution of the species to the community over a longer time (i.e. 50-200 generations). Overall, we hypothesize that the presence of the competitor can alter any evolutionary (genotype sorting to adaptive) response. For example, for *E. huxleyi*, the presence of the diatom assimilating CO_2_ could alleviate CO_2_ stress, and thus reduce selection towards tolerance, slowing down its adaptation in co-culture.

## 2 Material and Methods

We exposed two important phytoplankton species; the coccolithophore *E. huxleyi* and the diatom *C. affinis* to increased CO_2_ (ambient, high) either alone or in co-culture (mono, mix) with the respective other species over 288 days (=36 batch cycles of 8 days), representing approximately 200 asexual generations. We followed absolute species biomass at the end of each batch cycle over time to understand how the species respond to the different treatments. Each species started as a mixture of nine different genotypes to allow for rapid adaptation via genotype selection. All genotypes used were isolates originating from one geographical region (Gran Canary, 27°59’N 15°22’W, isolated 2012 and 2015) and are deposited in the culture collection in Roscoff (Table 1). In previous studies we have shown that the genotypes differed in growth, and to some extent in their response to CO_2_ (Hattich et al., 2017) indicating ecological variability and as such standing genetic variation. After 64 days (ca. 50 generations) and 288 days (ca. 200 generations), we conducted reciprocal adaptation experiments to test for adaptation to CO_2_ with and without competition. Ideally, to fully disentangle selection to inter- and intraspecific competition and the intended abiotic treatment single genotype cultures under both CO_2_ conditions would be required. However, this was technically not feasible for the phytoplankton species in this study as it would have required to handle another 180 bottle’s each batch cycle.

**Table 1:**
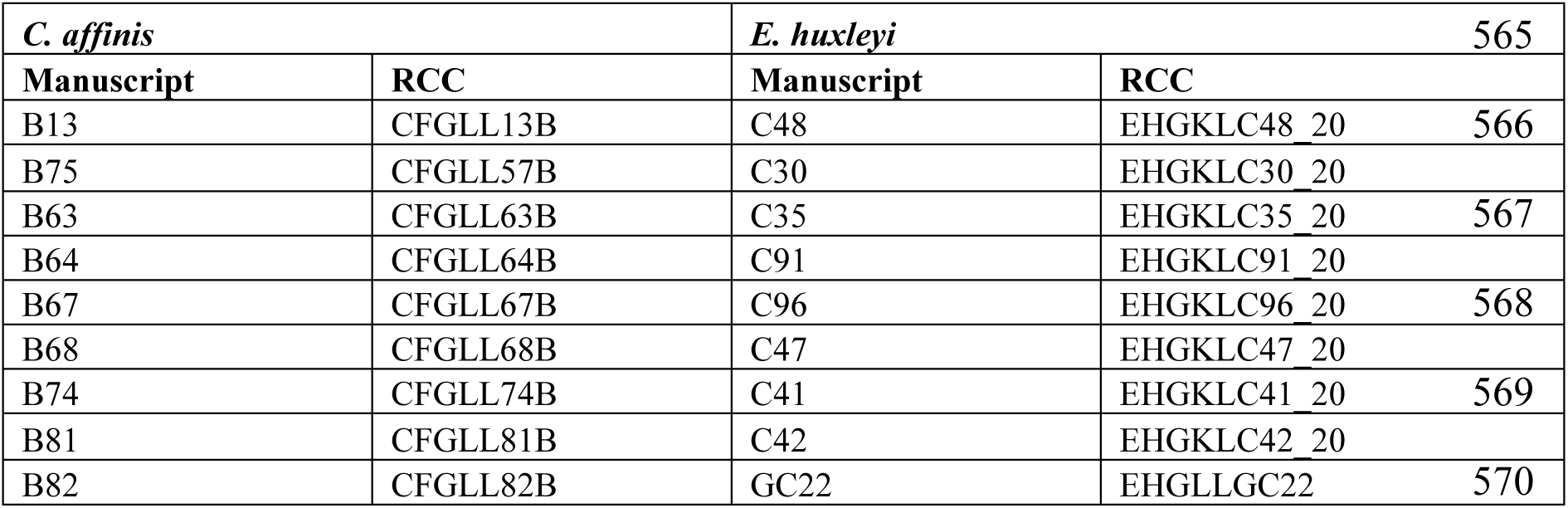
All genotypes used are listed in the column “manuscript” under the name used in here and are deposited in the Roscoff Culture Collection (RCC) under the name shown in the “RCC” column.

### Experimental set up and culturing conditions

The selection treatments were set-up in a fully orthogonal way with the single factors CO_2_ (two levels, ambient and high) and co-occurrence or absence of a 2^nd^ species (Mono and Mix) resulting in 4 treatment combinations per species (Fig. 1 a). Five replicate cultures of each of the four treatment combinations were kept for 9 months (36 batch cycles, resulting in 210-230 generations in *E. huxleyi* and 195-210 generations in *C. affinis*) in semi-continuous batch cultures. However, after ca. 28-30 batch cycles one replicate culture per treatment combination was lost in *E. huxleyi* and one in 3 out of 4 treatment combinations in *C. affinis*. The ambient and high CO_2_-treatments were manipulated by aerating the artificial-seawater (35 PSU salinity; after (Kester et al., 1967)) for 24h with ambient or CO_2_-enriched air (400 and 1250ppm, respectively) prior to two batch cycles. The dissolved inorganic carbon (Hansen et al., 2013) varied between 2052.39±34.49 and 2202±8.19 μmol*kg^-1^ with a total alkalinity (following (Matthiessen et al., 2012)) of 2272.56±11.24 and 2273.12±11.43 μmol*kg^-1^ for ambient and high CO_2_, respectively. Nutrients were added to the final concentrations of 19.61±0.09 μmol*L^-1^ nitrate, 0.99±0.02 μmol*L^-1^ phosphate and 4.34±0.14 μmol*L^-1^ silicate. In order to simulate a bloom situation with resource competition under depleted nutrients (Paulton, 1991), we ran the study as semi-continuous batch cultures with both species reaching the stationary phase (supplementary Figure S1 growth graphs of the different species). This resulted in the almost full depletion of all macronutrients (nitrate, phosphate and silicate) at the end of the batch cycles (supplementary Fig S2). Each experimental unit was inoculated with an initial total biomass of 8280 μm^3^*mL^−1^. When the two species were growing together, they were added at a ratio of 1:1 of the biomass to start the experiment. The nutrient concentrations and starting conditions were determined in prior pilot studies (Listmann and Hattich unpublished data) to ensure coexistence for at least 50-100 generations. The experiment was carried out under constant rotation (0.75 min^−1^) at ∼22 °C and a 17:7 day:night cycle reaching a maximum light intensity of 350 μmol*m^-2^*s^-1^ three hours after dusk and dawn. At the end of each batch cycle cell numbers and cell volume were determined using microscopy (Zeiss Axiovert Observer) to calculate the contribution of each species to the total biomass and the amount needed to transfer to the next batch cycle; again 8280 μm^3^*mL^-1^.

### Frequency assessments of genotypes

In order to follow genotype frequencies, we re-isolated cells of both species after 8, 32, 64, 160 and 288 days. Genotypes of both species could be unambiguously identified via microsatellite genotyping (see (Hattich et al., 2017), for detailed information on *E. huxleyi* and for *C. affinis* genotype identification see supplementary material). For the quantification of *E. huxleyi* and *C. affinis* genotypes a maximum of 20 cells per culture were re-isolated by dilution in 48 well plates. This provided a theoretical detection limit of 5% difference between the contributions of the genotypes to both species’ populations. A lower detection limit could in theory have been achieved via isolation and identification of more cells but was not the goal of this study. Details on the reisolation and quantification via microsatellite analysis are given in supplementary material. For the genotype composition of *C. affinis* we only had enough data for analysis two time points in the mixed cultures.

### Reciprocal adaptation assays

We carried out reciprocal adaptation assays to test for adaptation in both species to enhanced CO_2_, the competition of the respective other species, and both factors in combination (Fig. 1 b). The assays on day 64 and 288 assessed how adaptation played out over time. Those adaptation assays compared evolved populations in control and new environments rather than evolved and ancestral populations because methods such as cryopreservation are not readily available for our target study species (Collins, 2011b). Specifically, every evolved culture was tested in all four treatment combinations in a full factorial way. For example, *E. huxleyi* that were long-term treated with high CO_2_ conditions (treatment combination Mono High), were in the assay exposed to both ambient and high CO_2_ concentrations and then also to the co-occurring diatom (already long-term exposed to ambient and high CO_2_ and competition to avoid confounding responses) in ambient and high CO_2_ concentrations. Thus, each treatment combination from the selection leads to 4 assay treatment combinations. This resulted in a total of 16 assay treatment combinations per species (Fig. 1 b; for the details of the reciprocal assay set up see supplementary material). Using a full factorial adaptation assay allows us to test on the one hand for the adaptation of the single factor CO_2_ (i.e. in the statistical analysis, this could be identified as significant interactions of the respective selection and assay treatment factors (e.g. selection CO_2_ x assay CO_2_)). On the other hand, we can additionally test for the adaption to increased CO_2_ in combination with competition but then also how competition itself affected adaptive responses. This allows disentangling the single and combined treatment factors at the same time. Here we focused on growth rates because this response is directly related to Darwinian fitness in an asexual population (Elena and Lenski, 2003) and is independent of nutrient availability due to the presence of another species (Tilman, 1977).

### Statistical analyses

In all treatments with co-occurring species, the relative species contribution of *E. huxleyi* to total biomass (Mix) over time was analyzed using a generalized least squares (GLS) model (m0<-gls (relative Biomass ∼ Selection CO_2_ * Time)). Differences in variance structure were adjusted for the factor “Selection CO_2_”, and accounted for autocorrelation over time. For the statistical analysis it was sufficient to only look at the relative contribution of one of the two species, as their respective contribution to the community was complementary. For the analysis of the absolute biomass of each species in all cultures we used the same GLS model. The analysis was done for each species separately and started with the following full model: m0<-gls (Biomass ∼ Selection CO_2_ * Selection Culture * Time). After reduction and analysis of the model structure we had to account for autocorrelation over time and differences in variance structure (varIDENT) for the “Selection Culture” in *E. huxleyi* and “Selection CO_2_” and “Selection Culture” in *C. affinis*. There was a strong change around 160 days of experiment and we found significant interactions with “Time” for all factors in the GLS model. In order to investigate the effects of the different treatments on both species before and after this time point, we divided the time series data into two parts – BC1-BC20 and BC21-BC36 on which we repeated the described GLS model (*BC1-20* and *BC21-36*, respectively<-gls (Biomass ∼ Selection CO_2_ * Selection Culture)).

A permutational multivariate analysis of variance (permanova, with 999 permutations (using the package “vegan”)) was used to test for the differences in genotype compositional change between the treatments.

To analyse the reciprocal assay data we used repeated measure analysis of variance (rmANOVA). Since during the experiment, 4 and 3 cultures were lost, we had to omit the data of the “lost” replicates from all statistical analyses of the assays. Before starting the analyses we tested for the homogeneity of variance using a Fligner Killeen Test (Fligner and Killeen, 1976) and for the normality of the residuals using a Shapiro-Wilk Normality Test (Shapiro and Wilk, 1965). The assumption of sphericity was not violated because we only had one repeated measure (Field et al.). All assumptions were met such that we could continue with the analysis without data transformation. The growth rate was first assessed for the overall effects of “Time” (between the two assays), “CO_2_” and “Culture” in both selection and assay environment and thus we started with a complete data set analysis (aov((muExp)∼(Selection CO_2_ * Selection Culture * Assay CO_2_ * Assay Culture * Time) + Error(Replicate/Time))) followed by a separate analysis of each assay. Here, interactions of selection treatment * assay treatment would indicate evolutionary adaptation over the course of the experiment.

All modelling and statistical analyses were done using the software R (Coreteam, 2016). The following packages were used for the analyses, plotting and modelling: ’ggplot2’, ’deSolve’, ’vegan’, ’ez’ (Michael A. Lawrence, 2016; Oksanen et al., 2007; Soetaert et al., 2010; Wickham, 2009).

## 3 Results

### Changes in species composition over time

The outcome of interspecific competition in ambient and high CO_2_ was mirrored in relative biomass shifts: In the first half of the experiment the diatom dominated the two-species community with a relative biomass (mean over first 20 batch cycles) of ca. 95% in high compared to ambient CO_2_ condition with 80% (Fig. 2, Table S4 *BC1-20* “Selection CO_2_” F_1,186_=164.039, p<0.0001). From ca. 160 days onwards there was not only a dominance reversal from *C. affinis* to *E. huxleyi* (Fig. 2, Table S4 *Full Model* “Time” F_1,337_=613.093, p<0.0001, “Selection CO_2_*Time” F_1,337_=26.036, p<0.0001), but also a different reaction of both species to CO_2_, with *E. huxleyi* being favored by high CO_2_ in the second phase of the experiment. The reversal of dominance was reflected in an almost significantly decreased final relative contribution of *C. affinis* of 43% and 37% in ambient and high CO_2_ conditions, respectively (Fig. 2, Table S4 *BC21-36* “Selection CO_2_” F_1,142_=3.266, p=0.072).

**Figure 2.**
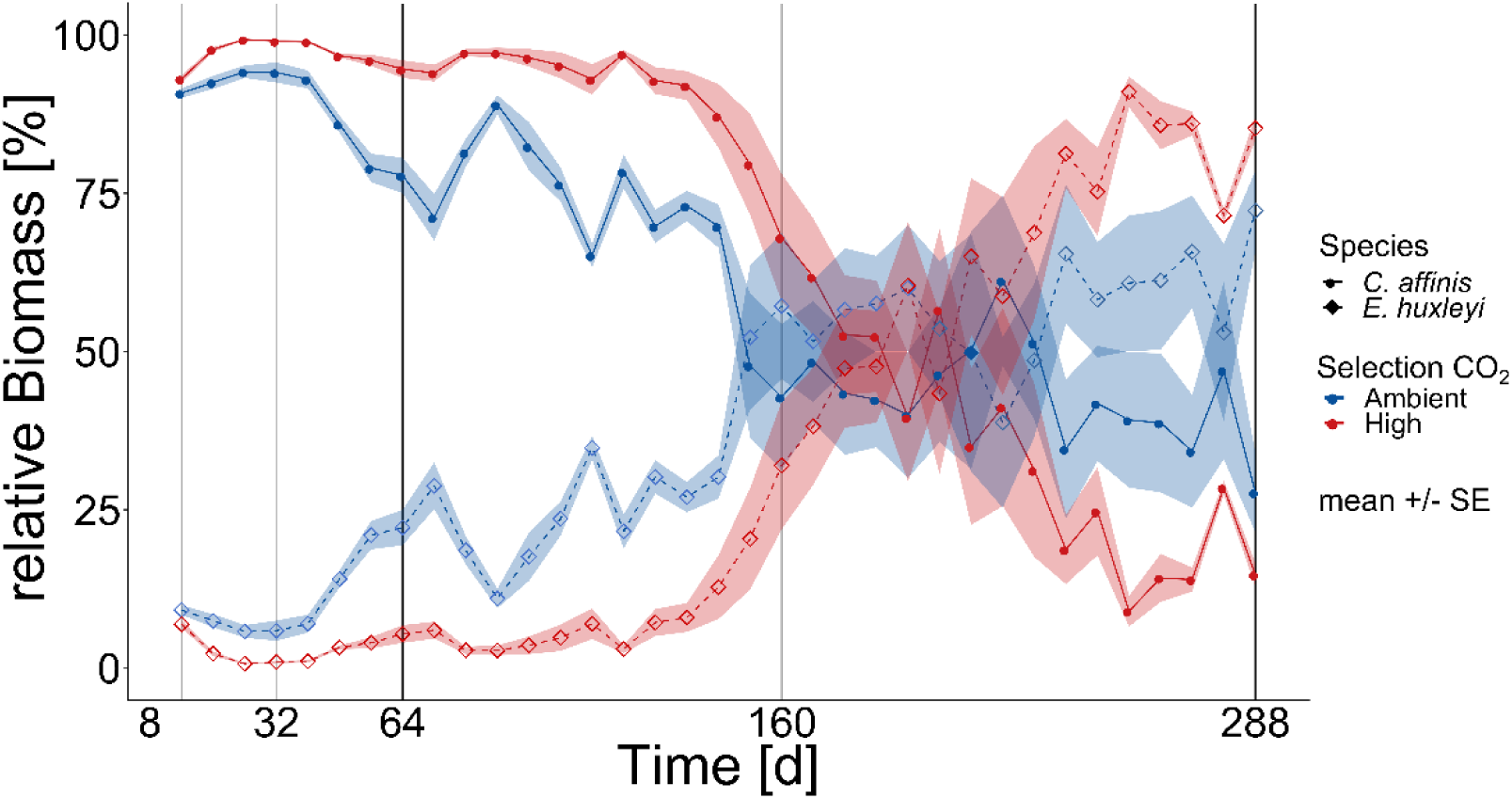
Community biomass composition. The relative contribution based on biomass of *E. huxleyi* and *C. affinis* measured at the end of each batch cycle over 36 batch cycles in the mix cultures (i.e. co-occurrence of both species) in ambient and high CO_2_ is shown here (mean ±SE; n=5for days 1-200 and n=4 for days 200-288 per time point). The light and dark grey lines show the time points where samples were taken for genotype reisolation. The dark grey lines show when the cultures were additionally tested for adaptation in the reciprocal assay.

### Temporal dynamics of absolute species biomass

Similar to the dynamics of relative species contributions, the time course of absolute biomasses of both species was characterized by two distinct phases that changed around 160 days of the experiment (Fig. 3 a-d, Table S5 *E. huxleyi* and *C. affinis* single and interactive effects of “Time” in *Full model*).

**Figure 3.**
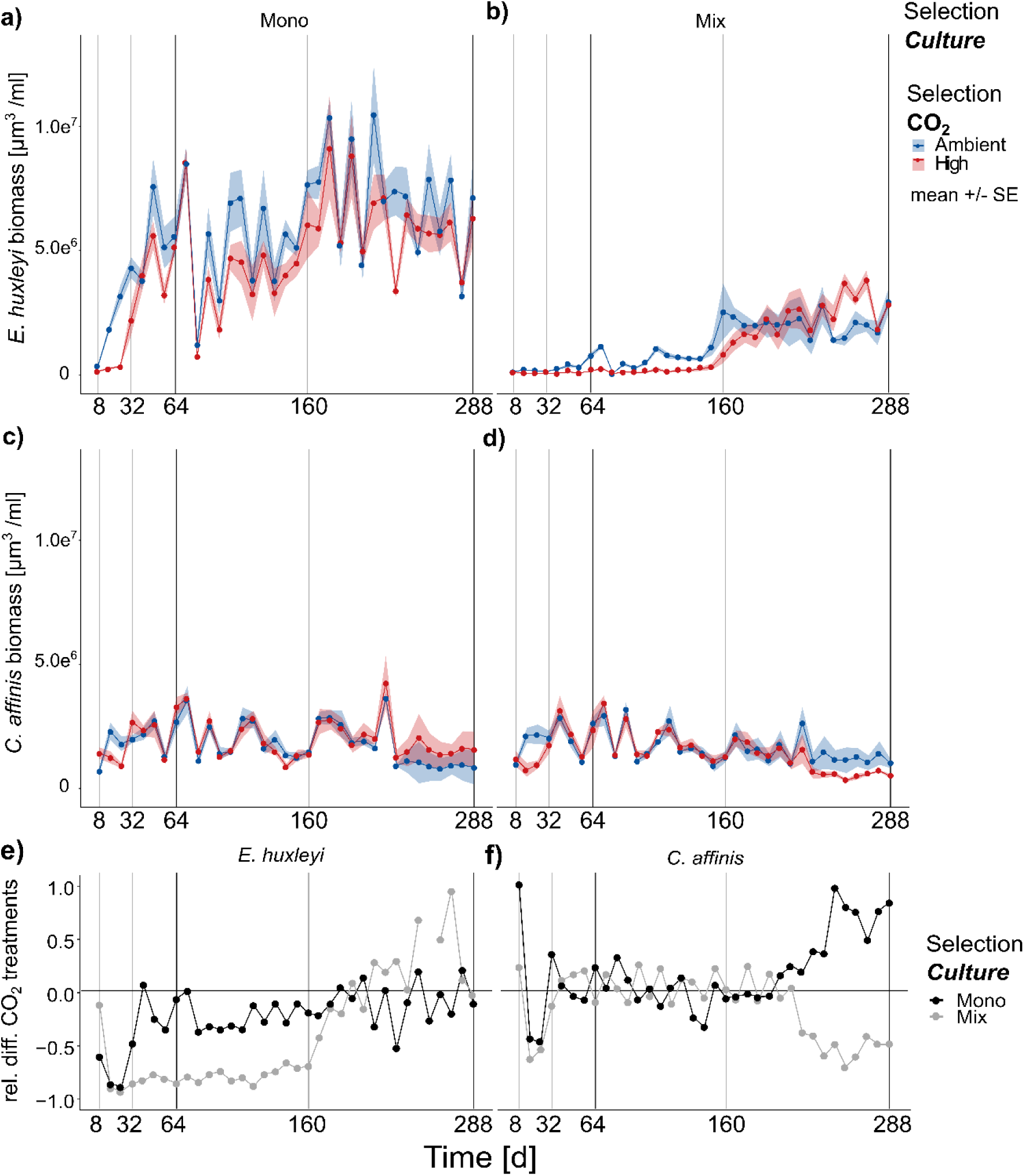
Biomass changes over approx. 200 generations. The biomass of *E. huxleyi* (**a, b**) and *C. affinis* (**c, d**) measured at the end of each batch cycle over 36 batch cycles in the four selection treatments is shown here (mean ±SE; n=5 (5 replicates/treatment) for days 1-200 and n=4 (4 replicates/treatment) for days 200-288 per time point). The light grey and black lines show the time points where samples were taken for genotype reisolation. The black lines show when the cultures were additionally tested for adaptation in the reciprocal assay test. Panels **e)** and **f)** show the relative difference between the ambient and high CO_2_ treatments for *E. huxleyi* (e) and *C. affinis* (f) when grown alone (black points) and with the respective other species (grey points) for better understanding how the responses to increased CO_2_ concentrations changed over time.

Overall *E. huxleyi* biomass increased over time about one order of magnitude with an average 30-fold increase across all treatments (Fig 3a and b, Table S5 *Full model*, “Time” F_1,660_=122.726, p<0.0001). During the first half of the experiment high CO_2_ reduced the biomass of *E. huxleyi* by a third in the mono-cultures (Fig. 3a and e Table S5 *E. huxleyi BC1-20* “Selection CO_2_” F_1,372_=113.818, p<0.0001). The effect of high CO_2_ was, with a biomass reduction of almost 4 times compared to the ambient treatment, even more negative on *E. huxleyi* in mix-cultures (Fig. 3b and e, Table S5 *E. huxleyi BC1-20* “Selection CO_2_*Selection Culture” F_1,372_=8.313, p=0.0042). This suggested that interspecific competition had an additive negative effect at first. In the second half of the experiment the effect of competition reversed and *E. huxleyi* biomass was only slightly reduced in high CO_2_ in the mono-cultures whereas in mix-culture increased CO_2_ even had a slightly positive effect (Fig. 3a, b and e, Table S5 *E. huxleyi BC21-36* “Selection CO_2_*Selection Culture” F_1,280_=7.156, p=0.0079).

In contrast to *E. huxleyi* the biomass of *C. affinis* varied markedly between batch cycles with only a slight decrease of biomass over time (Fig 3c and d, Table S5 *C. affinis Full model*, “Time” F_1,671_=25.538, p<0.0001). Specifically, the biomass of *C. affinis* was not affected by increased CO_2_ in the first half of the experiment, neither in mix- or mono-culture (Fig. 3c and d, Table S5 *C. affinis BC1-20*, “Selection CO_2_” F_1,372_=0.534, p=0.465). This pattern changed towards the end of the experiment where *C. affinis* biomass declined to half in the high CO_2_ treatment in the mix-cultures (Fig. 3d and f) while it doubled in the mono-cultures (Fig. 3c and f, Table S5 *C. affinis BC21-36*, “Selection CO_2_ * Selection Culture” F_1,352_=15.986, p<0.0001). This suggests that, over time, the diatoms in mix-culture were affected by the increasing biomass of the coccolithophores leading to diverging responses to high CO_2_ concentrations with and without competition (Table S5 *C. affinis Full model*, three-way interaction F_1,671_=12.225, p=0.0005).

### Genotype sorting

The genotype composition changed uniformely over time in both species (Fig. 4, Table S6, *E. huxleyi* “Time” R^2^=0.999, p=0.001, permutations=999, *C. affinis* “Time” R^2^=0.999, p=0.001, permutations=999) with only a single genotype left for all cultures of *E. huxleyi* and *C. affinis* (C41 and B57, respectively). In *E. huxleyi* there was no effect of CO_2_ on the genotype sorting (Fig. 4, Table S6, “Selection CO_2_” R^2^=0, p=0.401, permutations=999) and only a small difference between the single and co-occurring cultures (Fig. 4, Table S6 “Selection culture” R^2^=0, p=0.085, permutations=999). The small difference likely came about due to slightly slower sorting to the dominant genotype C41 in the mix-culture (Fig. 4 a vs b). For *C. affinis* we could not analyze the effect of either treatment on the genotype composition.

**Figure 4.**
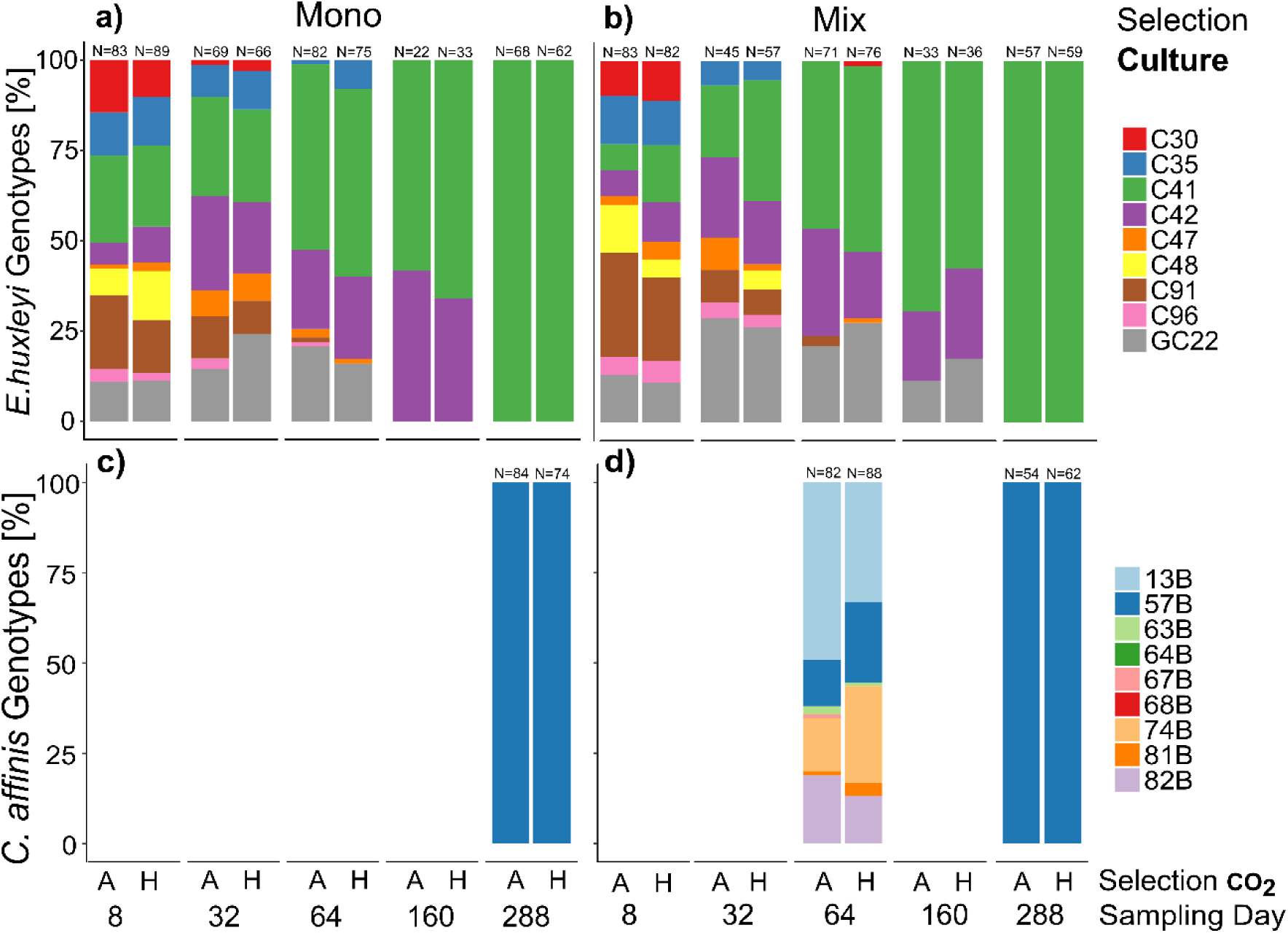
Genotype sorting over approx. 200 generations. Relative genotype contributions in *E. huxleyi* (**a** and **b**) and *C. affinis* (**c** and **d**) experimental populations at 5 and 2 timepoints, respectively, over the course of the experiment in ambient (A) and high (H) CO_2_ concentrations. Both panels on the left show the genotype contribution to the population without the second species (mono-culture) whereas the panels on the right show the contribution to the population with the respective other species (mix-culture). Re-isolation and DNA extraction of *C. affinis* genotypes proved difficult and we did not obtain usable data. However, the overall picture of strong genotype sorting remained.

### Reciprocal adaptation experiments

In *E. huxleyi* we found no significant interaction of selection with the assay treatment in neither of the two assays which would indicate evolutionary adaptation (Table S7, *E. huxleyi*). Specifically, long-term selection to high CO_2_ did not result in increased growth in the high CO_2_ assay condition compared to ambient selected populations (Fig 5a and b). However, there were non-adaptive effects of CO_2_ and Culture selection and assay treatments that varied between the two assay experiments: After 64 days, selection under high CO_2_ had a negative effect on the growth rates of *E. huxleyi* when measured in the presence of the diatom (“mix” assay treatment), while “mix” assay conditions led to a decline in growth rates in general (Fig. 5a and b, Table S7 *E. huxleyi Assay 64 days* “Selection CO_2_ * Assay Culture” F_1,47_=10.511, p=0.002, “Assay Culture” F_1,47_=254.708, p<0.0001). These effects were largely absent in the second assay after 288 days where growth rates were only affected by the assay CO_2_ treatment (Table S7 *E. huxleyi Assay 64 days*, F_1,47_= 8.066, p= 0.007). There was a significant increase in growth rates of ca. 10-45% over time in all selection treatments (Fig 5a; Table S7 *E. huxleyi*, “Time” F_1,47_=391.511, p<0.0001), with the strongest increase in cultures that had been selected in high CO_2_ in the presence of the diatom.

**Figure 5.**
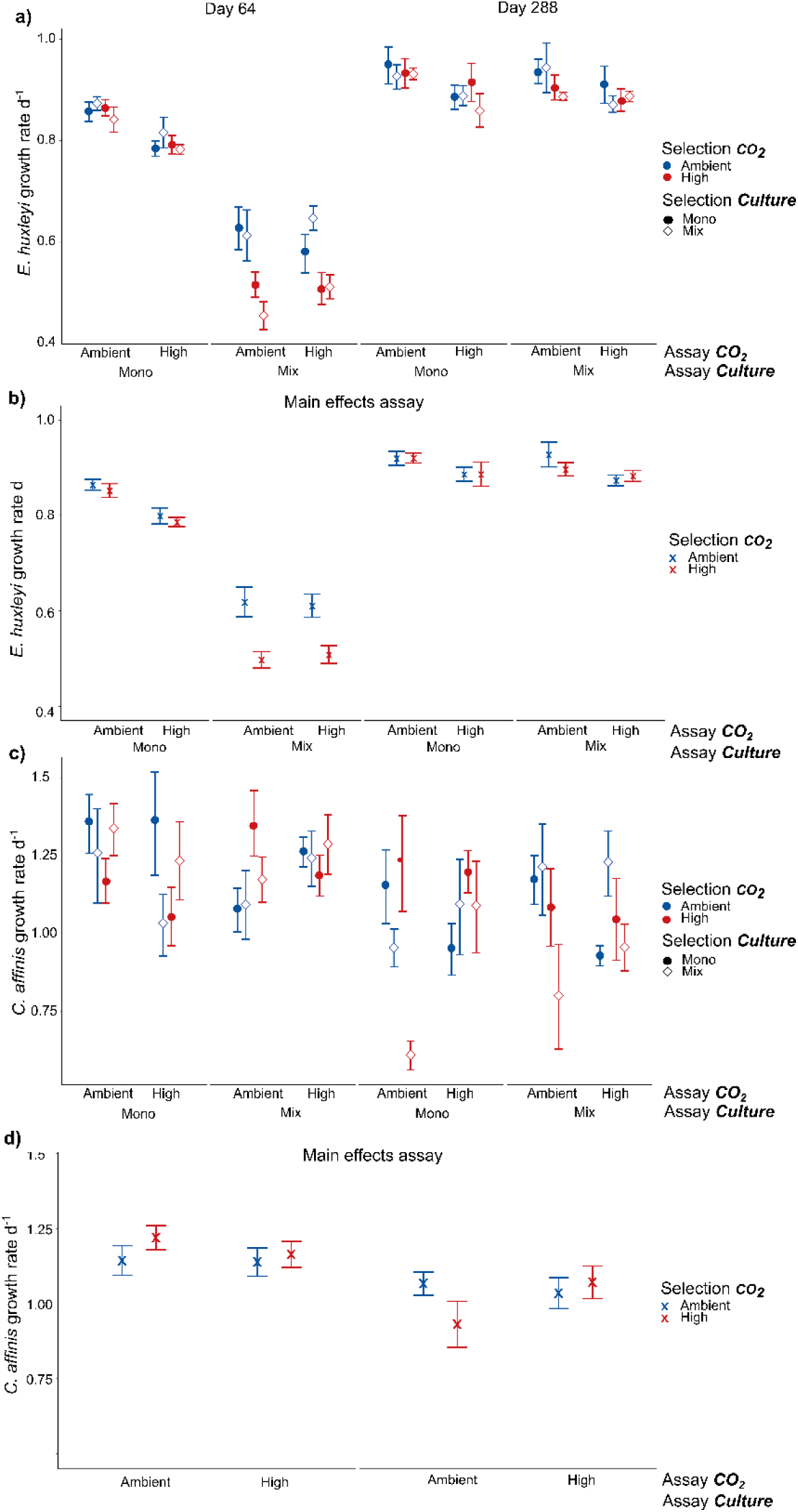
Growth rates in reciprocal assay experiments. Panels **a** and **c)** show the full results of adaptation assay experiments on growth rates of *E. huxleyi* and *C. affinis*, respectively after 64 and 288 days (mean ±SE, n=4). Panels **b** and **d)** show the significant and interactive results of the assays where we pooled the growth rates of the mono and mix selection culture treatments for both *E. huxleyi* and *C. affinis* and additionally the assay mono and mix culture treatment for *C. affinis* because these treatments did not have a significant effect (mean ±SE, n=16 and n=8, respectively). How to look for adaptive response: To see an adaptive response to the single factor CO_2_ it is necessary to compare all points with closed circles in Mono. For the adaptation to CO_2_ in a community context all points with open diamonds in Mix have to be compared. For the combined effect of adaption to CO_2_ and competition the comparison of the red diamond and blue circle in Mix-High and Mono-Ambient have to be compared.

No selection for higher growth rates to exposure to increased CO_2_ could be observed in *C. affinis*. In contrast to this expectation the growth rates decreased by a tenth over time throughout all treatments (Fig. 5 c, Table S7 *C. affinis* “Time” F_1,46_=14.843, p<0.001). Nevertheless, we found different responses to CO_2_ with and without competition that even resulted in an seemingly adaptive effect to increased CO_2_ in the second assay after 288 days, evident as a significant interaction of selection and assay treatment (Fig 5 d “u-shape” pattern in pooled data, Table S7 *C. affinis Assay 288 days* “Selection CO_2_” * “Assay CO_2_” F_1,47_=17.807, p=0.0001). However, this “adaptive” response was mainly driven by very low growth rates of *C. affinis* that have been selected in mix-cultures to high CO_2_ as well as low growth rates of *C. affinis* selected under ambient CO_2_ in mono-cultures. This result showed that disentangling the effects of both abiotic and biotic factors is crucial to avoid overestimation of adaptive responses to abiotic factors alone.

## 4 Discussion

Our results show that long-term selection in communities to an abiotic stressor can drastically diverge from the predicted outcomes of short term experiments with the same abiotic driver (Hattich et al., 2017; Riebesell, 2004). This divergence potentially arises from the effect of evolutionary on ecological processes, as previously shown in other systems (Hairston et al., 2005; Schoener, 2011). During the first half of the experiment our initial expectations (for biomass) with respect to species being an ecological “loser” and “winner” in response to enhanced CO_2_ were met for *E. huxleyi* but not *C. affinis*.

This led to the expected dominance of *C. affinis*, which was more pronounced under increased CO_2_ conditions. However, over time, the observed effects of increased CO_2_ on biomass changed for both species so as to revert our predicted ecological “loser” and “winner” outcome. Consequently, the relative contributions in the communities flipped and *E. huxleyi* became dominant under both CO_2_ conditions. We note that the “dominance-flip” coincided with strong genotype selection in both species, which was however not driven by CO_2_ or competition. Likewise, neither species showed adaptation to CO_2_ or competition in the reciprocal assays.

The absent selective and adaptive response to CO_2_ in both species, is for coccolithophores in contrast to previous studies (Lohbeck et al., 2012) and for diatoms both in contrast (Scheinin et al., 2015) and agreement (Li et al., 2017). The contrasting finding of no adaption or differential genotype sorting to increased CO_2_ for *E. huxleyi* could potentially be explained by differences in the experimental setups concerning the CO_2_ manipulation, batch cycle length, experimental duration, light, temperature and nutrient availability (Gao and Campbell, 2014; Meyer and Riebesell, 2015; Rost et al., 2008). A marked difference of our experimental setup compared to other experimental evolution studies was that our species were not kept under constant exponential growth (Li et al., 2017; Lohbeck et al., 2012; Pardew et al., 2018) but ran into stationary phase. The resulting recurrent nutrient limitation could likely override CO_2_ effects, as observed herin (or f.e. in Aranguren-Gassis, Kremer, Klausmeier, & Litchman, 2019). For *C. affinis*, however, it remains uncertain if nutrient limitation has likewise overwritten the expected adaptive response to CO_2_ or if the species was really unaffected by this driver. There is no directly comparable study with the same species, which does not run into nutrient limitation. Moreover the two long-term studies existing for diatoms do show diverging adaptive responses, which might depend on different experimental set-ups as well as species identity, as largely diverging short-term responses have already been described between diatom species (Bach and Taucher, 2019; Gao and Campbell, 2014).

It is well accepted that a more complex ecological context allowing for competition can affect evolutionary dynamics (Barraclough, 2015; de Mazancourt et al., 2008; Lancaster et al., 2017) but we did not find a significant effect in this study. Although not demonstrated to date, it is quite likely that interspecific competition has the potential to affect evolution in phytoplankton, as already intraspecific competition was shown to alter adaptive responses to CO_2_ (Collins, 2011a). Here, the dominant *C. affinis* strongly reduced the biomass and growth rate of *E. huxleyi*, demonstrating that the effect of competition was strong, especially in the first half of the experiment. In addition, we found a further reduction of growth rates to increased CO_2_ with the co-occurrence of the diatom indicating that the ecological context did play a role albeit not affecting the evolutionary change. That no effect of competition on evolution, or adaptation to competition could be observed potentially depends on how coexistence was allowed in our experiment. Owing to the species’ different nutrient uptake strategies (Sommer, 1984) niche partitioning likely appeared temporary over the course of a batch cycle. Whereas the diatom had an advantage during the first batch cycle days with replete nutrients, the coccolithophore was favored and could grow longer towards the end of each batch cycle when nutrients became limiting (Sommer, 1984) (Fig. S1-1 and S1-2). As such selection on standing genotype composition likely took place within these two niche spaces of the both coexisting species, unaffected by an interspecific competitor.

Despite the absence of an adaptive response to increased CO_2_ or competition, we still observed strong evolutionary change as very reproducible, directional sorting of genotypes across all treatments. Interestingly, these evolutionary dynamics selected for one single genotype of each species that became dominant already after approximately 64-160 days of the experiment. While we know that evolution was independent of the intended CO_2_ and competition treatments, non-intended laboratory selection was the most likely driver in our study. Laboratory selection is inevitable and often strong and has been previously demonstrated over very long time scales by comparing ancestral bacterial populations to laboratory evolved populations (Lenski and Travisano, 1994). Here, selection became apparent in an overall increased *E. huxleyi* and decreased *C. affinis* biomass and changing growth rates over time (Fig. S1-3). Interestingly, an increase of *E. huxleyi* growth rate was also observed in another long-term laboratory selection study (Schlüter et al., 2016). In these experiments however, laboratory adaptation was only a “background” signal, whereas in our experiments the chosen treatments were too weak to impose a selection force to overcome laboratory selection. Schlüter et al (2016) started their experiment with one single genotype, implying that observed changes in growth rate were a result of novel mutations, while in our study, observed changes in biomass and growth rate could be the result of both selection on genotypic diversity and mutations. Among the single remaining genotype it is possible that within the 200 generations of experimentation new mutations occurred (Elena and Lenski, 2003) that may contribute to phenotypic divergence in genotypes solely characterized by microsatellite marker alleles. With the caveat of not having ancestral populations of genotypes as a comparison we cannot say how laboratory selection played out in our species. Certain is however, that changes in species’ characteristic as observed in changed biomass and growth rate responses to the treatments throughout the experiment were strong and could have far reaching effects on ecosystem dynamics.

When studying eco-evolutionary dynamics, complex ecological conditions can evidently impinge upon evolutionary dynamics, while the complementary question asks how evolutionary change affects ecological processes (for example Ellner et al., 2011; Fussmann et al., 2007; Koch et al., 2014). In our study, the genotype compositional change (i.e. strong evolutionary change) coincided or rather preceded reversal in community composition (i.e. ecological change), suggesting a feedback from evolutionary changes on ecology in all treatments. That this dominance shift and underlying genotype selection also occurred in an earlier attempt of the experiment, suggests that the eco-evolutionary dynamics observed are reproducible (Fig. S1-4 and S1-5). We propose that selection by nutrient limitation on standing variability in genotype’s competitive abilities was the most likely driver resulting in the eco-evolutionary interaction. However, necessary experimental characterization of the genotypes’ nutrient uptake associated traits is (currently) missing. Other studies reported nutrient uptake related traits in marine protists to vary between genotypes to the same extent as growth rates (Brandenburg et al., 2018). Since we have shown that the variability in growth rates among the genotypes used in this study is even larger than that of the response to increased CO_2_ (Hattich et al., 2017), we postulate that their nutrient uptake related traits vary as well. Another indication that nutrients were driving the change in our system is the observed shift away from *C. affinis* towards *E. huxleyi*, the favored species by nutrient limitation (Tyrrell and Merico, 2004). Consequently, future studies should focus more on the consequences on how nutrients select to predict phytoplankton change (Thomas et al., 2017). This will become particularly relevant as nutrient uptake related traits explain competitive ability under different nutrient conditions (Litchman et al., 2007; Sommer, 1984) and there will be more nutrient limitation on phytoplankton in the future ocean (Boyd et al., 2013; Moore et al., 2013).

To conclude, the most parsimonious explanation for the observed dominance switch among the two globally important phytoplankton species *E. huxleyi* and *C. affinis*, is rapid evolutionary feedback back to the ecology. Rapid evolutionary change as genotype sorting driven by experimental selection and not CO_2_ environment turned an ecological “loser” (with respect to biomass contribution to the community) into a “winner” and vice versa. As such, this flip demonstrates for the first time that eco-evolutionary interactions play out in competing phytoplankton communities. Such interactions can drastically alter the effect of environmental drivers and lead to diverging predictions of future changes compared to such resulting of short-term studies. Our results call for an inclusion of more realistic experimental evolution conditions in future studies, not only using realistic nutrient regimes, but more importantly also including multi-species settings and their underlying mechanisms allowing for stable coexistence to simultaneously investigate ecological and evolutionary processes in phytoplankton.

## 5 Acknowledgements

We thank Thomas Hansen, Bente Gardeler, Cordula Meyer, Jens Wemhöner, Kastriot Qelai for their laboratory assistance. As well as our student assistants Nele Rex, Julia Raab, Julia Romberg, Miriam Beck, Sophia Antoniella and Gabriela Escobar for their support. We thank Till Bayer for his support with the development of *C. affinis* microsatellite assays. We also thank KIMOCC for technical support and quality management of the Kiel CO2 manipulation experimental facility (KICO2) at GEOMAR. We thank Prof. Dr. Elisa Schaum for her comments to improve this manuscript.

## 6 Author Contributions

The basic grant proposal supporting this work was written by BM and TBHR. The experiment was developed by GISH and then the experiments further designed by LL and GISH. Lab work carried out by LL and GISH and data analyses by LL. LL, GISH, BM and TBHR drafted the manuscript and all authors revised the manuscript and gave final approval for publication.

## 9 Data availability statement

All data will be made available through PANGEA upon publication. https://doi.pangaea.de/10.1594/PANGAEA.895771

## 10 Supplementary Material

Further methodological explanations on reciprocal assay experiments and genotype identification

***Table S4-S7***: statistical analysis of the selection experiment, reciprocal adaptation assays and genotype sorting respectively

***Figure S1***: Semi continuous batch cycle growth curves

***Figure S2***: Dissolved inorganic nutrients in one batch cycle

***Figure S3***: Estimated growth rates over the course of the experiment

***Figure S4***: relative species contribution to biomass from a first attempt at the experiment presented here

***Figure S5***: relative genotype contribution to biomass from a first attempt at the experiment presented here

## References

Aranguren-Gassis, M., Kremer, C. T., Klausmeier, C. A., and Litchman, E. (2019). Nitrogen limitation inhibits marine diatom adaptation to high temperatures. Ecol. Lett. 22, 1860–1869. doi: 10.1111/ele.13378.

Bach, L., and Taucher, J. (2019). CO_2_ effects on diatoms: A Synthesis of more than a decade of ocean acidification experiments with natural communities. Ocean Sci. Discuss., 1–40. doi: 10.5194/os-2019-47.

Barraclough, T. G. (2015). How Do Species Interactions Affect Evolutionary Dynamics Across Whole Communities? Annu. Rev. Ecol. Evol. Syst 46, 25–48. doi: 10.1146/annurev-ecolsys-112414-054030.

Boyd, P. W., Rynearson, T. A., Armstrong, E. A., Fu, F., Hayashi, K., Hu, Z., et al. (2013). Marine phytoplankton temperature versus growth responses from polar to tropical waters--outcome of a scientific community-wide study. PLoS One 8, e63091. doi: 10.1371/journal.pone.0063091.

Brandenburg, K. M., Wohlrab, S., John, U., Kremp, A., Jerney, J., Krock, B., et al. (2018). Intraspecific trait variation and trade-offs within and across populations of a toxic dinoflagellate. Ecol. Lett. 21, 1561–1571. doi: 10.1111/ele.13138.

Carroll, S. P., Hendry, A. P., Reznick, D. N., and Fox, C. W. (2007). Evolution on ecological timescales. Funct. Ecol. 21, 387–393. doi: 10.1111/j.1365-2435.2007.01289.x.

Collins, S. (2011a). Competition limits adaptation and productivity in a photosynthetic alga at elevated CO2. Proc. Biol. Sci. 278, 247–55. doi: 10.1098/rspb.2010.1173.

Collins, S. (2011b). Many Possible Worlds: Expanding the Ecological Scenarios in Experimental Evolution. Evol. Biol. 38, 3–14. doi: 10.1007/s11692-010-9106-3.

Coreteam, R. (2016). R: A Language and Environment for Statistical Computing.

de Mazancourt, C., Johnson, E., and Barraclough, T. G. (2008). Biodiversity inhibits species’ evolutionary responses to changing environments. Ecol. Lett. 11, 380–8. doi: 10.1111/j.1461-0248.2008.01152.x.

Des Roches, S., Post, D. M., Turley, N. E., Bailey, J. K., Hendry, A. P., Kinnison, M. T., et al. (2018). The ecological importance of intraspecific variation. Nat. Ecol. Evol. 2, 57–64. doi: 10.1038/s41559-017-0402-5.

Doney, S. C., Fabry, V. J., Feely, R. A., and Kleypas, J. A. (2009). Ocean Acidification : The Other CO2 Problem. 169–194.

Elena, S. F., and Lenski, R. E. (2003). Evolution experiments with microorganisms: The dynamics and genetic bases of adaptation. Nat. Rev. Genet. doi: 10.1038/nrg1088.

Ellner, S. P., Geber, M. a, and Hairston, N. G. (2011). Does rapid evolution matter? Measuring the rate of contemporary evolution and its impacts on ecological dynamics. Ecol. Lett. 14, 603–14. doi: 10.1111/j.1461-0248.2011.01616.x.

Falkowski, P. G., Fenchel, T., and Delong, E. F. (2008). The Microbial Engines That Drive Earth ‘s Biogeochemical Cycles. Science (80-.). 320, 1034–1039. doi: 10.1126/science.1153213.

Field, A., Field, A., and |α, S. A Bluffer’s Guide to… Sphericity. Available at: http://citeseerx.ist.psu.edu/viewdoc/summary?doi=10.1.1.161.3892 [Accessed April 11, 2018].

Fligner, M. A., and Killeen, T. J. (1976). Distribution-Free Two-Sample Tests for Scale. J. Am. Stat. Assoc. 71, 210–213. doi: 10.1080/01621459.1976.10481517.

Fussmann, G. F., Loreau, M., and Abrams, P. a. (2007). Eco-evolutionary dynamics of communities and ecosystems. Funct. Ecol. 21, 465–477. doi: 10.1111/j.1365-2435.2007.01275.x.

Gao, K., and Campbell, D. A. (2014). Photophysiological responses of marine diatoms to elevated CO 2 and decreased pH: a review.

Hairston, N. G., Ellner, S. P., Geber, M. A., Yoshida, T., and Fox, J. A. (2005). Rapid evolution and the convergence of ecological and evolutionary time. Ecol. Lett. 8, 1114–1127. doi: 10.1111/j.1461-0248.2005.00812.x.

Hansen, T., Gardeler, B., and Matthiessen, B. (2013). Technical Note: Precise quantitative measurements of total dissolved inorganic carbon from small amounts of seawater using a gas chromatographic system. Biogeosciences 10, 6601–6608. doi: 10.5194/bg-10-6601-2013.

Hardin Garret (1960). The Competitive Exclusion Principle. Science (80-.). 131, 1292–1297. Available at: https://www.jstor.org/stable/1705965.

Hattich, G. S. I., Listmann, L., Raab, J., Ozod-Seradj, D., Reusch, T. B. H., and Matthiessen, B. (2017). Inter-and intraspecific phenotypic plasticity of three phytoplankton species in response to ocean acidification. Biol. Lett. 13. doi: 10.1098/rsbl.2016.0774.

Hendry, A. P. (2016). Eco-evolutionary dynamics. Princeton university press.

Kester, D. R., Duedall, I. W., Connors, D. N., and Pytkowicz, R. M. (1967). Preparation of artificial seawarter. Limnol. Oceanogr. 12, 176–179. doi: 10.4319/lo.1967.12.1.0176.

Koch, H., Frickel, J., Valiadi, M., and Becks, L. (2014). Why rapid, adaptive evolution matters for community dynamics. Front. Ecol. Evol. 2, 1–10. doi: 10.3389/fevo.2014.00017.

Lancaster, L. T., Morrison, G., and Fitt, R. N. (2017). Life history trade-offs, the intensity of competition, and coexistence in novel and evolving communities under climate change. Philos. Trans. R. Soc. B Biol. Sci. 372, 20160046. doi: 10.1098/rstb.2016.0046.

Lawrence, D., Fiegna, F., Behrends, V., Bundy, J. G., Phillimore, A. B., Bell, T., et al. (2012). Species interactions alter evolutionary responses to a novel environment. PLoS Biol. 10, e1001330. doi: 10.1371/journal.pbio.1001330.

Lenski, R. E., and Travisano, M. (1994). Dynamics of adaptation and diversification: a 10,000-generation experiment with bacterial populations. Proc. Natl. Acad. Sci. U. S. A. 91, 6808–14. doi: 10.1073/PNAS.91.15.6808.

Li, F., Beardall, J., Collins, S., and Gao, K. (2017). Decreased photosynthesis and growth with reduced respiration in the model diatom Phaeodactylum tricornutum grown under elevated CO2 over 1800 generations. Glob. Chang. Biol. 23, 127–137. doi: 10.1111/gcb.13501.

Litchman, E., Klausmeier, C. a., Schofield, O. M., and Falkowski, P. G. (2007). The role of functional traits and trade-offs in structuring phytoplankton communities: Scaling from cellular to ecosystem level. Ecol. Lett. 10, 1170–1181. doi: 10.1111/j.1461-0248.2007.01117.x.

Lohbeck, K. T., Riebesell, U., and Reusch, T. B. H. (2012). Adaptive evolution of a key phytoplankton species to ocean acidification. Nat. Geosci. 5, 346–351. doi: 10.1038/ngeo1441.

Matthiessen, B., Eggers, S. L., and Krug, S. A. (2012). High nitrate to phosphorus regime attenuates negative effects of rising pCO 2 on total population carbon accumulation. Biogeosciences 95194, 1195–1203. doi: 10.5194/bg-9-1195-2012.

Meyer, J., and Riebesell, U. (2015). Reviews and Syntheses : Responses of coccolithophores to ocean acidification : a meta-analysis. 1671–1682. doi: 10.5194/bg-12-1671-2015.

Michael A. Lawrence (2016). ez: Easy Analysis and Visualization of Factorial Experiments. Available at: https://cran.r-project.org/package=ez.

Moore, C. M., Mills, M. M., Arrigo, K. R., Berman-Frank, I., Bopp, L., Boyd, P. W., et al. (2013). Processes and patterns of oceanic nutrient limitation. Nat. Geosci. 6. doi: 10.1038/ngeo1765.

Oksanen, J., Kindt, R., Legendre, P., O ‘hara, B., Henry, M., and Maintainer, H. S. (2007). The vegan Package Title Community Ecology Package. Available at: http://ftp.unibayreuth.de/math/statlib/R/CRAN/doc/packages/vegan.pdf [Accessed April 17, 2018].

Pardew, J., Blanco Pimentel, M., and Low-Decarie, E. (2018). Predictable ecological response to rising CO 2 of a community of marine phytoplankton. Ecol. Evol.

Paulton, R. J. L. (1991). The bacterial growth curve. J. Biol. Educ. 25, 92–94. doi: 10.1080/00219266.1991.9655183.

Reusch, T. B. H., and Boyd, P. W. (2013). Experimental Evolution Meets Marine Phytoplankton. Evolution (N. Y). 67, 1849–1859. doi: 10.1111/evo.12035.

Reznick, D. N. (2013). A Critical Look at Reciprocity in Ecology and Evolution: Introduction to the Symposium. Am. Nat. 181, S1–S8. doi: 10.1086/670030.

Riebesell, U. (2004). Effects of CO2 enrichment on marine phytoplankton. J. Oceanogr. 60, 719–729. doi: 10.1007/s10872-004-5764-z.

Rost, B., Zondervan, I., and Wolf-Gladrow, D. (2008). Sensitivity of phytoplankton to future changes in ocean carbonate chemistry: Current knowledge, contradictions and research directions. Mar. Ecol. Prog. Ser. 373, 227–237. doi: 10.3354/meps07776.

Schaum, E., Rost, B., Millar, A. J., and Collins, S. (2012). Variation in plastic responses of a globally distributed picoplankton species to ocean acidification. Nat. Clim. Chang. 3, 298–302. doi: 10.1038/nclimate1774.

Scheinin, M., Riebesell, U., Rynearson, T. a, Lohbeck, K. T., and Collins, S. (2015). Experimental evolution gone wild. J. R. Soc. Interface 12, 1–5. doi: 10.1098/rsif.2015.0056.

Schlüter, L., Lohbeck, K. T., Gröger, J. P., Riebesell, U., and Reusch, T. B. H. (2016). Long-term dynamics of adaptive evolution in a globally important phytoplankton species to ocean acidification. Sci. Adv. 2, e1501660. doi: 10.1126/sciadv.1501660.

Schoener, T. W. (2011). The newest synthesis: understanding the interplay of evolutionary and ecological dynamics. Science 331, 426–429. doi: 10.1126/science.1193954.

Shapiro, S. S., and Wilk, M. B. (1965). An Analysis of Variance Test for Normality (Complete Samples). Biometrika 52, 591. doi: 10.2307/2333709.

Soetaert, K., Petzoldt, T., and Setzer, R. W. (2010). Solving Differential Equations in R: Package deSolve. J. Stat. Softw. 33, 1–25. doi: 10.18637/jss.v033.i09.

Sommer, U. (1984). The paradox of the plankton: Fluctuations of phosphorus availability maintain diversity of phytoplankton in flow-through cultures. Limnol. Oceanogr. 29, 633–636. doi: 10.4319/lo.1984.29.3.0633.

Sommer, U., Paul, C., and Moustaka-Gouni, M. (2015). Warming and ocean acidification effects on phytoplankton - From species shifts to size shifts within species in a mesocosm experiment. PLoS One 10, 1–17. doi: 10.1371/journal.pone.0125239.

Thomas, M. K., Aranguren-Gassis, M., Kremer, C. T., Gould, M. R., Anderson, K., Klausmeier, C. A., et al. (2017). Temperature–nutrient interactions exacerbate sensitivity to warming in phytoplankton. Glob. Chang. Biol. 23, 3269–3280. doi: 10.1111/gcb.13641.

Tilman, D. (1977). Resource Competition between Plankton Algae : An Experimental and Theoretical Approach Author (s): David Tilman Published by : Wiley on behalf of the Ecological Society of America Stable URL : http://www.jstor.org/stable/1935608 REFERENCES Linked refere. 58, 338–348.

Tyrrell, T., and Merico, A. (2004). Emiliania huxleyi: bloom observations and the conditions that induce them. Coccolithophores, 75–97. doi: 10.1007/978-3-662-06278-4_4.

Wickham, H. (2009). ggplot2: Elegant Graphics for Data Analysis. Springer-Verlag New York. Available at: http://ggplot2.org.

