## Supplementary Material for "Eco-evolutionary interaction in competing phytoplankton: genotype sorting likely explains dominance shift and species responses to CO_2_"

### Supplementary material to Manuscript Listmann et al.:

### Additional Material and Methods Descriptions:

#### Genotype quantification via microsatellite analysis:

For the quantification of *E. huxleyi* genotypes a maximum of 20 cells per culture were re-isolated by dilution in 48 well plates. This provides a theoretical detection limit of 5% difference between the contributions of the genotypes to the *E. huxleyi* population. When left unmoved in culture plates, a single *E. huxleyi* cell forms a bacterial-like "benthic" colony, which can be easily detected after ten days. Fresh medium (500µL) was added to each well with growing colonies and cells were grown for another ten days. Cells were then centrifuged and the pellets of *E. huxleyi* were used for DNA extraction and further microsatellite analysis. *E. huxleyi* DNA was extracted by adding 15µL of TE-buffer to re-suspend the pellet. Samples were then sonicated for three minutes at 100% and then incubated for 1h at 56°C. Microsatellite amplification was done following the protocol described in (Hattich et al., 2017) with the following primers: Primer S15 (Access. No. AJ487032 (F) AJ487305 (R)) and Primer E10 (Access. No. AJ487314 (F) AJ487315 (R)). The relative contribution of specific genotypes to the total population of *E. huxleyi* was calculated as (no. of certain genotype/no. of genotypes analyzed per culture). In *C. affinis* we re-isolated a minimum of 20 cells by dilution into 48 well plates. Where microscopy (magnification 10x) had revealed the presence of a single cell/chain on the first day after reisolation, after 8 days the cultures were transferred to new medium to grow for another 8 days. The cells were then centrifuged, and the DNA of the resulting pellet was extracted using the Quiagen® DNeasy Blood & Tissue 96-well plate kit. We found that very small amounts of material (<1ng/µl) were not enough to yield adequate microsatellite analysis results (no usable data). Microsatellite amplification was done following the protocol described in supplementary material with the following primers: Primer C.a._LL_8 and Primer C.a._LL_17 (Primer sequences in Table S1).

#### ***Chaetocers affinis*** **microsatellite assays**

*DNA extraction*

DNA from two strains of *C. affinis* was extracted using a modified CTAB extraction method (Gagnon et al., 1980). The samples were frozen in liquid nitrogen and crushed with a cold adjusted pistil. Then CTAB extraction buffer as well as 100µL of RNAse were added followed by an incubation for 15 minutes at 60°C and 15 minutes at room temperature. Two steps of elution with a chloroform-isoamyl alcohol mixture (24:1 vol:vol) were done. The DNA was the precipitated from the water phase with ice cold isopropanol for 30 minutes. The DNA was additionally cleaned with 70% EtOH. With this method high quality DNA containing long strands can be extracted. The yields were 19.1 ng/µL for G05786 (strain B81) and 52.2 ng/µL for G05787 (strain B64).

*DNA sequencing and primer search and testing*

Sequencing libraries with 100 bp insert size were generated from genomic DNA using the TruSeq DNA Nano protocol. The libraries were sequenced on an Illumina NextSeq500, resulting in 33.5 and 17.5 million read pairs of 150 bp length for the samples G05786 and G05786, respectively. Raw reads were quality trimmed to an average quality of 10 and adapters clipped using BBDuk from the BBTools suite (Bushnell) and then assembled into one draft genome per strain using the Platanus software with default parameters (Kajitani et al., 2014). Suitable microsatellite loci were searched in the resulting scaffolds using Tandem Repeats Finder (Benson, 1999), and primers for amplification were designed with the help of the Primer3 software (Untergasser et al., 2012, 3). Microsatellite loci with different alleles between the two strains of each species were selected for analysis.

The primers (Table S1) were tested on the nine genotypes used in the experiment and yielded positive microsatellite amplification in most strains. The primers were tested in the following reaction: 2.5µl multiplex mastermix (Qiagen), 1µL forward and reverse Primer pair, 2µl Q solution (Qiagen), 1µL $H_2O$ and 1µL of DNA template. The PCR reaction run for microsatellite amplification was set up as follows: an initial phase of 15min at 95°C, 30 cycles of 30sec at 94°C, 90sec at 57°C and 1min at 72°C and final step of 30 min at 60°C. 1µL PCR product was then added to a mix of ROX and Hidi (Qiagen) of 0.25µL and 8.75µL, respectively and incubated for 3 min at 94°C to denature double stranded products. We employed capillary electrophoresis coupled to fragment analysis on an ABI 3130xl genetic analyzer to score the microsatellite composition using the software GeneMarker software. Primers C.a._LL_8 and C.a._LL_17 yielded the best allele combination results (Table S2.) and were used in the analysis of the genotype composition throughout the experiment.

**Table S1**: Primers for microsatellite analysis in *C. affinis*

| **Primer Name** | **Forward seq.** | **Reverse seq.** |
|---|---|---|
| C.a._LL_1 | ACTTACCATGACAACAGCAATG | ACCCATTGAGATTTGAGTTCATG |
| C.a._LL_3 | TGGTCATATGTCCTCTTTCGG | GGCAAACACAAACACAAACAC |
| C.a._LL_5 | CACTTTTGAAGGTACACAGTGG | ACGTTGGGAGAGCTATTGAG |
| ***C.a._LL_8*** | ***GACGCTGGTAGTTTCGTTTG*** | ***AGTCCCTTGGAAATGGACTG*** |
| C.a._LL_9 | GTGTAAGCGTAAAAGAATGCATG | AGTTCCTGCCTTCAGTCTTC |
| C.a._LL_11 | TGCATCTTGCTTGAGGAGG | AGTGCAAGGTTTACTCAATTCTG |
| C.a._LL_14 | TGAACCTGTGGATATGACGG | CACCCTTCTTTATATGATGAGCG |
| C.a._LL_15 | TCTGCTCTGCGTTGTAGTTC | CTTTGTGTTTGCCTCCTTCC |
| ***C.a._LL_17*** | ***GGGGTAATGAAATCTTTGGTGC*** | ***GTACTGATTATCAACAGGTGCTC*** |
| C.a._LL_19 | TAGGAATCATCGGCATCTGC | ATACGAAGGCCTTTCTGGAG |

**Table S2**: Allele results for all primers

| **Genotype** | **C.a._LL_F_1** | | | **C.a._LL_F_3** | | | **C.a._LL_F_5** | | **C.a._LL_F_8** | | **C.a._LL_F_9** | | |
|---|---|---|---|---|---|---|---|---|---|---|---|---|---|
| B13 | 185 | 185 | | 205 | 231 | | 218 | 218 | 219 | 231 | 214 | 218 | 226 |
| B57 | 163 | 165 | | 231 | 231 | | 218 | 218 | 217 | 221 | 226 | 226 | |
| B63 | 163 | 163 | | 231 | 231 | | 218 | 218 | 215 | 215 | 226 | 226 | |
| B64 | 0 | 0 | | 0 | 0 | | 212 | 212 | 0 | 0 | 218 | 226 | |
| B67 | 0 | 0 | | 157 | 231 | | 216 | 218 | 213 | 215 | 226 | 255 | |
| B68 | 79 | 99 | 145 | 231 | 231 | | 167 | 218 | 213 | 221 | 226 | 232 | |
| B74 | 0 | 0 | | 0 | 0 | | 213 | 218 | 221 | 231 | 226 | 232 | |
| B81 | 145 | 165 | | 157 | 205 | 231 | 167 | 218 | 217 | 217 | 232 | 255 | |
| B82 | 79 | 185 | | 157 | 231 | | 167 | 218 | 217 | 229 | 255 | 255 | |
| **Genotype** | **C.a._LL_F_11** | | | **C.a._LL_F_14** | | **C.a._LL_F_15** | | **C.a._LL_F_17** | | | **C.a._LL_F_19** | | |
| B13 | 78 | 98 | | 226 | 226 | 153 | 185 | 173 | 177 | | 0 | 0 | |
| B57 | 74 | 94 | | 226 | 226 | 163 | 163 | 177 | 181 | | 0 | 0 | |
| B63 | 74 | 94 | | 0 | 0 | 183 | 183 | 171 | 171 | | 174 | 174 | |
| B64 | 76 | 96 | | 0 | 0 | 0 | 0 | 177 | 181 | | 0 | 0 | |
| B67 | 0 | 0 | | 222 | 222 | 163 | 163 | 163 | 177 | | 174 | 178 | |
| B68 | 78 | 78 | | 222 | 231 | 153 | 163 | 167 | 181 | | 178 | 187 | |
| B74 | 74 | 96 | | 0 | 0 | 0 | 0 | 167 | 179 | | 0 | 0 | |
| B81 | 102 | 104 | | 228 | 228 | 153 | 183 | 163 | 177 | | 174 | 178 | |
| B82 | 102 | 104 | 262 | 224 | 224 | 171 | 183 | 169 | 177 | 181 | 178 | 187 | |

**Set up of Reciprocal Assay:**

As a prerequisite to understand the potential of adaptation of both species separately and to make the adequate species combinations in the reciprocal assay experiments, it was necessary to separate the diatom and coccolithophore by a 20µM mesh sieve. This was possible because they differed enough in size: the diameter of *E. huxleyi* was between 2.5 and 4µM whereas the length of *C. affinis* started at ca. 15µM. Additionally, *C. affinis* has long spines that make it bigger than the coccolithophore and kept it from going through the used sieve. The efficiency of this method had been established in pilot studies. In order to test for adaptation and not short-term acclimation we let the cultures acclimatize to the assay conditions for one entire batch cycle prior to measuring adaptation responses. The $CO_2$ treatment in the assays was manipulated as in the long-term experiment and was therefore a constant manipulated factor in the two assays. To test for selection *via* another species, the abundance of the co-occurring species was adjusted in the respective assays to represent the actual presence of the other species at the time point of adaptation testing and this varied among the two assays (Table S3 relative species contributions). Specifically, at the end of batch cycle 8 and 36, the relative contributions of each species to the mix cultures were determined in both $CO_2$ treatments. The mean of the relative composition was then used in the assays as the relative species composition in the respective "mix" culture treatments in both ambient and high $CO_2$ (Table S3 relative species contributions). The 2$^{nd}$ species we added in the assay treatments came from the same selection treatment as the assay treatment to avoid any confounding responses of the added species. Consequently, the conditions for the treatments "mix, ambient" and "mix, high" were not equal in relative strength in both assays. However, since the absolute biomass of *C. affinis* did not change over time the absolute competitive effect of *C. affinis* on *E. huxleyi* likely remained similar.

**Table S3**. Relative species contributions (mean±SE) to each co-evolving community at the end of batch cycle 8 and 36.

| **Treatment** | **Species** | **Batch cycle 8 [**% contribution] | **Batch cycle 36 [**% contribution] |
|---|---|---|---|
| "2$^{nd}$ species | *E. huxleyi* | 21.1±2.1 | 72.3±6.5 |
| | *C. affinis* | 78.9±2.1 | 27.7±6.5 |
| "$CO_2$, 2$^{nd}$ species" | *E. huxleyi* | 3.9±0.89 | 85.1±1.8 |
| | *C. affinis* | 96.1±0.89 | 14.9±1.8 |

#### Supplementary Tables:

**Table S4**: Analysis report for relative biomass of *E. huxleyi* in the two-species cultures: a GLS model was used. The models account for autocorrelation of the factor “$CO_2$”. A change in variance structure over time is assumed.

| ***Analysis relative species composition*** | ***Full model*** | | | ***BC1-20*** | | | ***BC21-36*** | | |
|---|---|---|---|---|---|---|---|---|---|
| ***Analysis species sorting*** | df | F-value | P-Value | df | F-value | P-Value | df | F-value | P-Value |
| **Selection $CO_2$** | **1** | **11.747** | **<0.0001** | **1** | **164.039** | **<0.0001** | 1 | 3.266 | 0.072 |
| ***Time*** | **1** | **613.0931** | **<0.0001** | **1** | **85.586** | **<0.0001** | **1** | **36.978** | **<0.0001** |
| ***Selection $CO_2$ × Time*** | **1** | **26.036** | **<0.0001** | **1** | **31.583** | **<0.0001** | 1 | 14.032 | 0.0003 |
| ***Residuals*** | **337** | | | **186** | | | 142 | | |

**Table S5**: Statistical analysis report for effects of $CO_2$, Culture, time and their respective interactions on absolute species biomass: a GLS model was used. The full model accounts for the entire experimental time, whereas BC 1-20 and BC 21-36 account for statistical testing of the experimental phases before and after the observed dominance shift, respectively. The model accounts for a difference in variance structure for *E. huxleyi* for the factor “Culture” whereas a difference in variance structure for the factor “$CO_2$” for and “Culture” *C. affinis* is assumed. Additionally, in *C. affinis* we accounted for autocorrelation over time.

| ***Emiliania huxleyi*** | ***Full model*** | | | ***BC1-20*** | | | ***BC21-36*** | | |
|---|---|---|---|---|---|---|---|---|---|
| ***Analysis species sorting*** | df | F-value | P-Value | df | F-value | P-Value | df | F-value | P-Value |
| **Selection $CO_2$** | **1** | **38.886** | **<0.0001** | **1** | **113.818** | **<0.0001** | 1 | 0.013 | 0.910 |
| ***Culture*** | **1** | **844.1817** | **<0.0001** | **1** | **370.927** | **<0.0001** | **1** | **378.149** | **<0.0001** |
| ***Time*** | **1** | **122.726** | **<0.0001** | **1** | **64.906** | **<0.0001** | 1 | 1.690 | 0.194 |
| ***Selection $CO_2$ × Selection Culture*** | **1** | **15.670** | **0.0001** | **1** | **8.313** | **0.0042** | **1** | **7.156** | **0.0079** |
| ***Selection $CO_2$ × Time*** | **1** | **4.442** | **0.0355** | **1** | **21.955** | **<0.0001** | **1** | **12.887** | **0.0004** |
| ***Selection Culture × Time*** | **1** | **6.6537** | **0.01** | **1** | **11.654** | **0.0007** | **1** | **12.213** | **0.0006** |
| *Selection $CO_2$ × Selection Culture × Time* | 1 | 0.163 | 0.686 | 1 | 0.394 | 0.530 | 1 | 0.944 | 0.332 |
| *Residuals* | 660 | | | 372 | | | 280 | | |

| ***Chaetoceros affinis*** | ***Full model*** | | | ***BC1-20*** | | | ***BC21-36*** | | |
|---|---|---|---|---|---|---|---|---|---|
| ***Analysis species sorting*** | df | F-value | P-Value | df | F-value | P-Value | df | F-value | P-Value |
| *Selection $CO_2$* | 1 | 0.0050 | 0.9434 | 1 | 0.534 | 0.465 | 1 | 0.379 | 0.539 |
| ***Selection Culture*** | **1** | **22.814** | **<.0001** | 1 | 1.811 | 0.179 | **1** | **26.231** | **<.0001** |
| ***Time*** | **1** | **25.538** | **<0.0001** | 1 | 0.853 | 0.356 | **1** | **30.622** | **<0.0001** |

| | | | | | | | | | |
|---|---|---|---|---|---|---|---|---|---|
| *Selection $CO_2$ × Selection Culture* | 1 | 9.677 | 0.0019 | 1 | 0.029 | 0.864 | **1** | **15.986** | **<0.0001** |
| *Selection $CO_2$ × Time* | 1 | 2.189 | 0.134 | 1 | 3.061 | 0.081 | 1 | 0.000 | 0.999 |
| *Selection Culture × Time* | 1 | 7.676 | 0.0058 | 1 | 0.061 | 0.803 | 1 | 2.934 | 0.089 |
| ***Selection $CO_2$ × Selection Culture × Time*** | **1** | **12.225** | **0.0005** | 1 | 2.733 | 0.099 | **1** | **14.203** | **0.0002** |
| *Residuals* | 671 | | | 372 | | | 352 | | |

**Table S6**: Permanova results for analysis of genotype compositional change; permutations=999

| ***Emiliania huxleyi*** | | | | | | ***Chaetoceros affinis*** | | | | | |
|---|---|---|---|---|---|---|---|---|---|---|---|
| ***Permanova*** | df | MS | F Model | $R^2$ | P-Value | ***Permanova*** | df | MS | F-value | $R^2$ | P-Value |
| ***Time*** | **4** | **92402** | **1855273** | **0.999** | **0.001** | ***Time*** | **1** | **84505** | **2883621** | **0.999** | **0.001** |
| *Selection Culture* | 1 | 0 | 4 | 0 | 0.085 | *Selection Culture* | 1 | 0 | 2 | 0 | 0.180 |
| *Selection $CO_2$* | 1 | 0 | 1 | 0 | 0.401 | *Time × Selection $CO_2$* | 1 | 0 | 2 | 0.0001 | 0.191 |
| *Time × Selection $CO_2$* | 4 | 0 | 1 | 0 | 0.324 | *Residual* | 16 | | | | |
| *Time × Selection Culture* | 4 | 0 | 1 | 0 | 0.346 | | | | | | |
| *Selection $CO_2$ × Selection Culture* | 1 | 0 | 0 | 0 | 1 | | | | | | |
| *Time × Selection $CO_2$ × Selection Culture* | 4 | 0 | 1 | 0.0001 | 0.452 | | | | | | |
| *Residual* | 80 | | | | | | | | | | |

**Table S7**: Analysis report for reciprocal adaptation assay (growth rate): a repeated measures ANOVA for comparison between assays at 64 and 288 days was used whereas factorial ANOVAS were used for the assays separately. Only the significant results are reported here.

| ***Emiliania huxleyi*** | | | | | ***Chaetoceros affinis*** | | | | |
|---|---|---|---|---|---|---|---|---|---|
| ***Analysis Assays after 64 and 288 days*** | df | MS | F-value | P-Value | ***Analysis Assays after 64 and 288 days*** | df | MS | F-value | P-Value |
| *Error: replicate* | | | | | *Error: replicate* | | | | |
| *Selection $CO_2$* | 1 | 0.0542 | 14.469 | <0.001 | none | | | | |
| *Assay $CO_2$* | 1 | 0.0386 | 10.311 | 0.002 | | | | | |

| | | | | | | | | | |
|---|---|---|---|---|---|---|---|---|---|
| *Assay Culture* | 1 | 0.5745 | 153.307 | <0.0001 | | | | | |
| *Selection $CO_2$× Assay Culture* | 1 | 0.0330 | | 0.005 | | | | | |
| *Assay $CO_2$× Assay Culture* | 1 | 0.0195 | | 0.027 | | | | | |
| *Residuals* | 47 | 0.0037 | | | *Error: replicate:Time* | | | | |
| *Error: replicate:Time* | | | | | Time | 1 | 0.6238 | 14.843 | <0.001 |
| *Time* | 1 | 1.4208 | 391.511 | <0.0001 | *Selection $CO_2$× Selection Culture × Time* | 1 | 0.4607 | 10.961 | 0.002 |
| *Selection $CO_2$× Time* | 1 | 0.0246 | 6.779 | 0.012 | *Selection $CO_2$× Assay $CO_2$× Time* | 1 | 0.2481 | 5.902 | 0.019 |
| *Assay Culture × Time* | 1 | 0.5125 | 141.223 | <0.0001 | *Selection $CO_2$× Assay Culture × Time* | 1 | 0.2967 | 7.061 | 0.011 |
| *Residual* | 47 | $3.921e^{12}$ | | | Residuals | 46 | 0.0420 | | |

| | | | | | | | | | |
|---|---|---|---|---|---|---|---|---|---|
| ***Assay 64 days*** | | | | | ***Assay 64 days*** | | | | |
| *Selection $CO_2$* | 1 | 0.0759 | 17.807 | 0.0001 | | | | | |
| *Assay Culture* | 1 | 1.0861 | 254.708 | <0.0001 | | | | | |
| *Selection $CO_2$× Assay Culture* | 1 | 0.0448 | 10.511 | 0.002 | | | | | |
| *Assay $CO_2$× Assay Culture* | 1 | 0.0277 | 6.485 | 0.014 | *Selection $CO_2$× Selection Culture × Assay Culture* | 1 | 0.19669 | 4.295 | 0.044 |
| *Residuals* | 47 | 0.0043 | | | Residuals | 48 | | | |

| | | | | | | | | | |
|---|---|---|---|---|---|---|---|---|---|
| ***Assay 288 days*** | | | | | ***Assay 288 days*** | | | | |
| | | | | | *Selection $CO_2$× Selection Culture* | 1 | 0.3318 | 6.808 | 0.012 |
| *Assay $CO_2$* | 1 | 0.025103 | 8.066 | 0.007 | *Selection $CO_2$× Assay $CO_2$* | 1 | 0.2844 | 5.834 | 0.020 |
| *Residuals* | 47 | 0.003112 | | | Residuals | 46 | 0.0487 | | |

### Supplementary Figures:

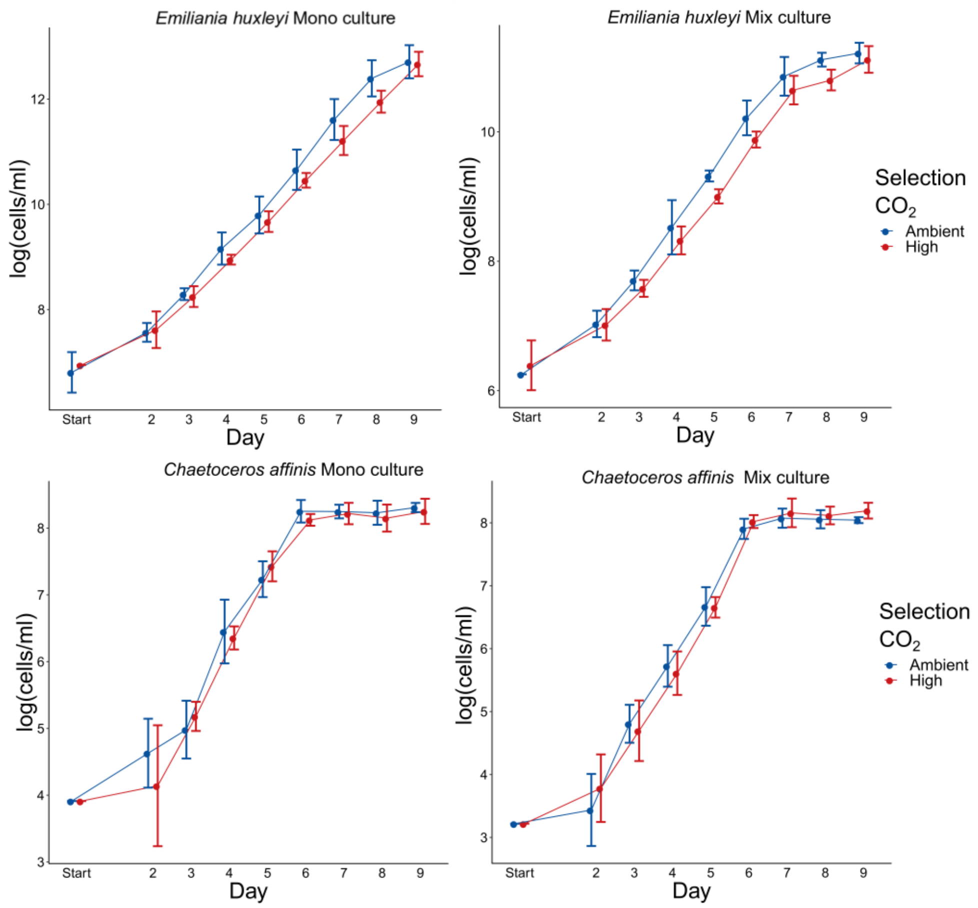


**Figure S1 Semi continuous batch cycle growth curve:** Here the growth within the first batch cycle is shown for each species and all four treatments used in the selection experiment. Log cell numbers highlight the exponential phase (linear increase) and stationary phase (flattening of the curve towards day 7-9). Each dot represents the mean ±SE, n=5.

*C. affinis* reached the stationary phase earlier than *E. huxleyi*, however both species reached the stationary phase in all the treatments; a requirement for competition in the system. The number of generations was calculated as gen=(ln($N_{max}$)-ln($N_0$))/$days_{growth}$ and ranged between 5-6 generations per batch cycle and species. As a diatom, *C. affinis* requires silicate to grow its silicate shells (Egge and Heimdal, 2012) and if silicate becomes limiting diatoms cannot grow anymore (Ragueneau et al., 2000). In order to ensure coexistence of the two species (unpublished data of preliminary experiments Listmann and Hattich) we set the silicate concentrations to ca 5µM in our study (see further details in methods).

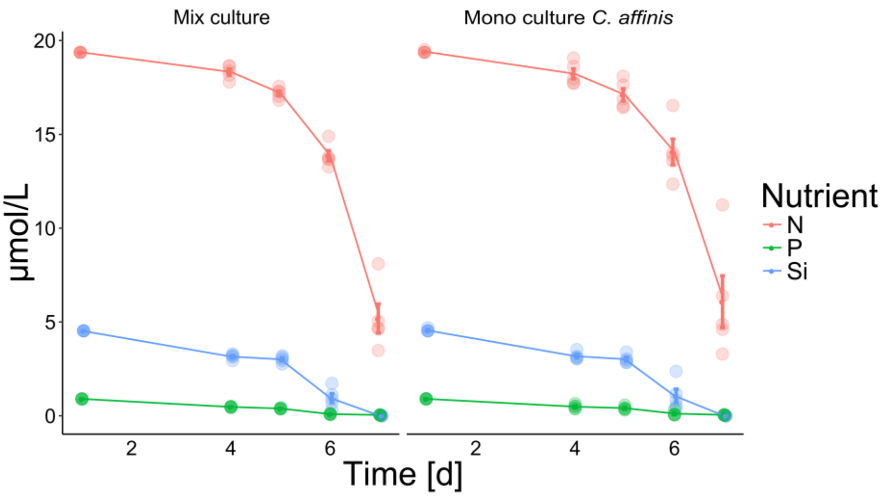


**Figure S2 Dissolved inorganic nutrients in one batch cycle**: Dissolved inorganic nutrients measured over one batch cycle in the cultures where *C. affinis* was present (BC11, day 40-48) (mean, n=4).

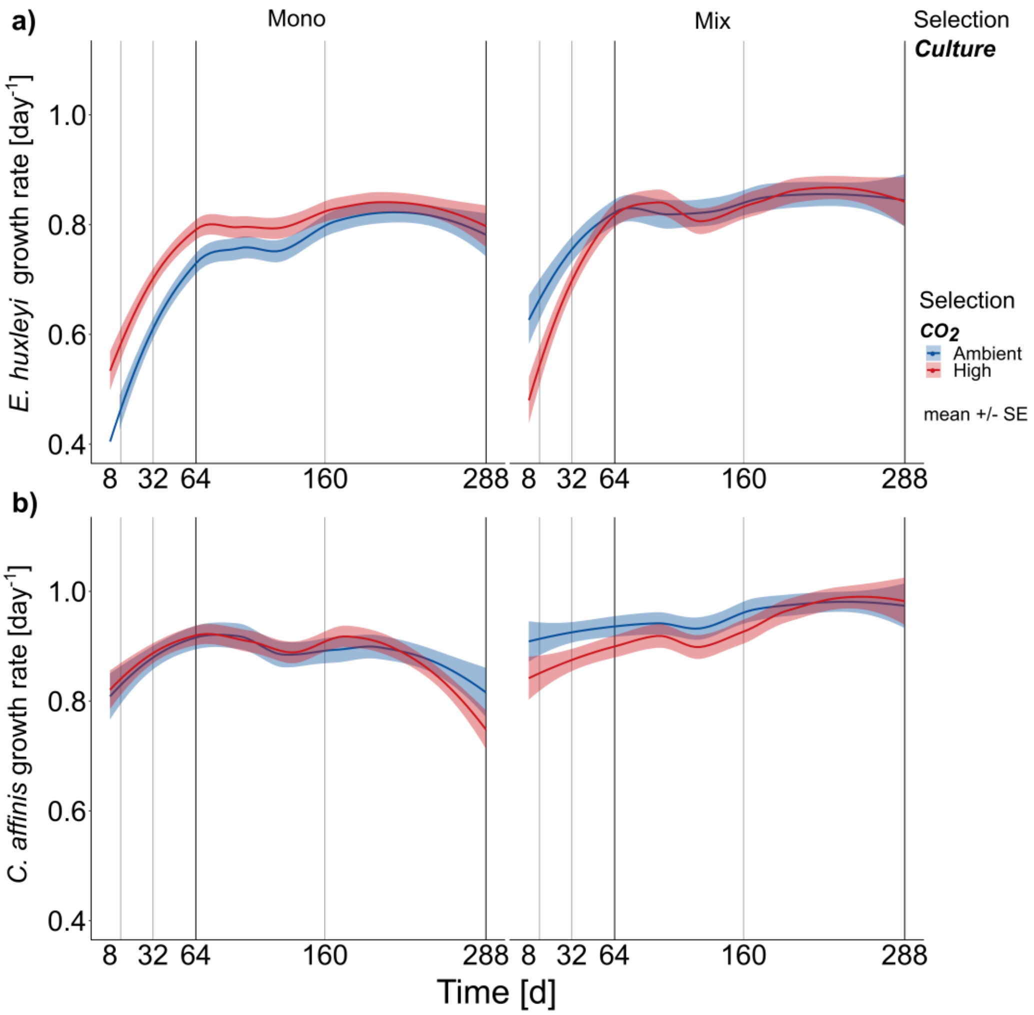


**Figure S3**. Estimated growth rates over the course of the experiment are shown here for E. huxleyi (top panel) and C. affinis (bottom panel). The growth rates were calculated based on start and end cell numbers of each batch cycle and an estimated length of the exponential growth days (8 and 6 for *E. huxleyi* and *C. affinis*, respectively). Shown here is the mean and smoother over the course of the experiment.

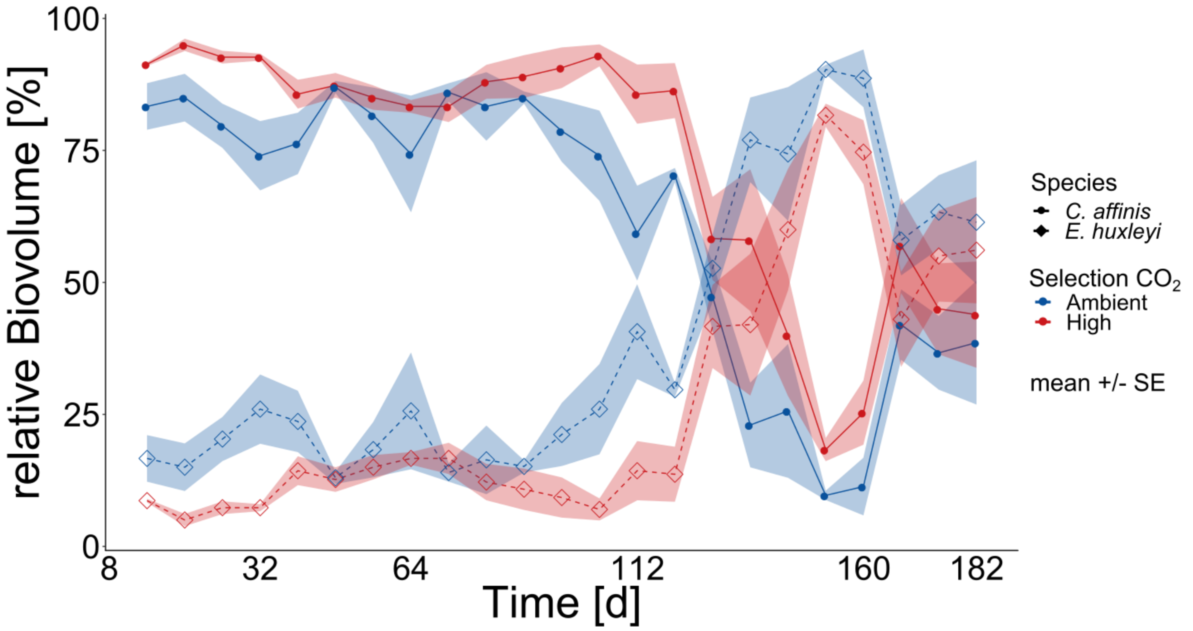


**Figure S4 relative species contribution to biomass from earlier experiment**. The relative contribution based on biomass of *E. huxleyi* and *C. affinis* measured at the end of each batch cycle over 24 batch cycles in the mix cultures in ambient and high $CO_2$ is shown here (mean ±SE; n=3 per treatment). The data were collected in a previous experiment with the same set up as described in the current study but had to be terminated owing to contamination and culturing problems between days 130 and 160.

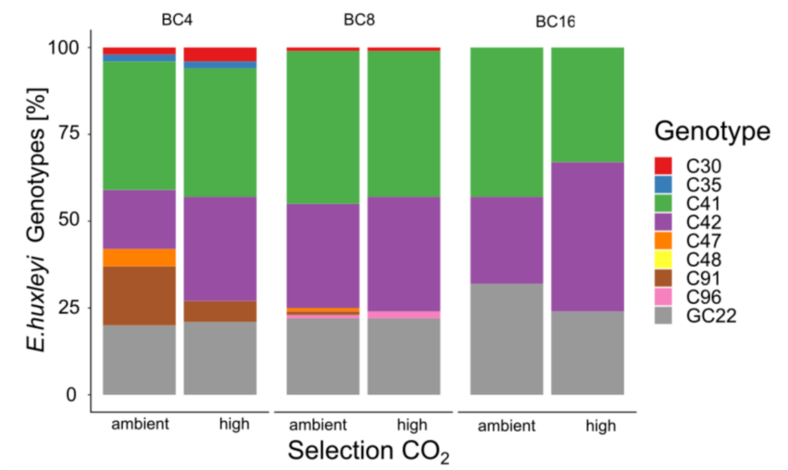


**Figure S5 relative genotype contribution from earlier experiment**. Relative genotype contributions in *E. huxleyi* experimental populations at 3 timepoints in, over the course of the experiment in ambient and high $CO_2$ concentrations. The data were collected in a previous experiment with the same set up as described in the current study but had to be terminated owing to contamination and culturing problems between days 130 and 160.

#### References

Benson, G. (1999). Tandem repeats finder: a program to analyze DNA sequences. *Nucleic Acids Res.* 27. Available at: http://citeseerx.ist.psu.edu/viewdoc/download?doi=10.1.1.69.9658&rep=rep1&type=pdf [Accessed March 19, 2018].

Bushnell, B. BBMap. *BBTools*.

Egge, J. K., and Heimdal, B. R. (2012). Blooms of phytoplankton including Emiliania huxleyi (Haptophyta). Effects of nutrient supply in different N : P ratios. *Sarsia*. Available at: http://www.tandfonline.com/doi/abs/10.1080/00364827.1994.10413565 [Accessed June 2, 2015].

Gagnon, P. S., Vadas, R. L., Burdick, D. B., and May, B. (1980). Genetic identity of annual and perennial forms of Zostera marina L. *Aquat. Bot.* 8, 157–162. doi:10.1016/0304-3770(80)90047-9.

Hattich, G. S. I., Listmann, L., Raab, J., Ozod-Seradj, D., Reusch, T. B. H., and Matthiessen, B. (2017). Inter-and intraspecific phenotypic plasticity of three phytoplankton species in response to ocean acidification. *Biol. Lett.* 13. doi:10.1098/rsbl.2016.0774.

Kajitani, R., Toshimoto, K., Noguchi, H., Toyoda, A., Ogura, Y., Okuno, M., et al. (2014). Efficient de novo assembly of highly heterozygous genomes from whole-genome shotgun short reads. *Genome Res.* 24, 1384–1395. doi:10.1101/gr.170720.113.

Ragueneau, O., Tréguer, P., Leynaert, A., Anderson, R. ., Brzezinski, M. ., DeMaster, D. ., et al. (2000). A review of the Si cycle in the modern ocean: recent progress and missing gaps in the application of biogenic opal as a paleoproductivity proxy. *Glob. Planet. Change* 26, 317–365. doi:10.1016/S0921-8181(00)00052-7.

Untergasser, A., Cutcutache, I., Koressaar, T., Ye, J., Faircloth, B. C., Remm, M., et al. (2012). Primer3—new capabilities and interfaces. *Nucleic Acids Res.* 40, e115–e115. doi:10.1093/nar/gks596.
